# Multi-marker metabarcoding resolves subtle variations in freshwater condition: Bioindicators, ecological traits, and trophic interactions

**DOI:** 10.1101/2021.11.14.468533

**Authors:** Chloe Victoria Robinson, Teresita M. Porter, Victoria Carley Maitland, Michael T.G. Wright, Mehrdad Hajibabaei

## Abstract

Freshwater systems are experiencing rapid biodiversity losses resulting from high rates of habitat degradation. Ecological condition is typically determined through identifying either macroinvertebrate or diatom bioindicator assemblages and comparing them to their known tolerance to stressors. These comparisons are typically conducted at family or genus levels depending on the availability of taxonomic keys and expertise for focal groups. The objective of this study was to test whether a more taxonomically comprehensive assessment of communities in benthic samples can provide a different perspective of ecological conditions. DNA metabarcoding was used to identify macroinvertebrates and diatoms from kick-net samples collected from sites with different habitat status. Sites with ‘good’ condition were associated with higher beta diversity as well as slightly higher directed connectance and modularity indicating higher resilience compared with ‘fair’ condition sites. Indicator value and correlation analyses used DNA metabarcoding data to detect 29 site condition indicator species consistent with known bioindicators and expected relative tolerances. DNA metabarcoding and trophic network analysis also recovered 11 keystone taxa. This study demonstrates the importance of taxonomic breadth across trophic levels for generating biotic data to study ecosystem status, with the potential to scale-up ecological assessments of freshwater condition, trophic stability, and resilience.

## Introduction

We are currently experiencing rapid freshwater biodiversity declines on a global scale, as anthropogenic pressures including habitat destruction, water pollution, overexploitation and climate change continue to escalate. The process of slowing future freshwater biodiversity losses is complicated, due to the influence of surrounding land use, particularly upstream human activities, on environmental conditions within rivers, lakes, wetlands and ponds ^1, 2^. Protecting and restoring aquatic ecosystems across spatial and temporal scales requires a multi-faceted approach, inclusive of ecological network analyses (e.g. trophic interactions) ^1, 3–5^. Before we can take the actions necessary to conserve freshwater ecosystems, we need to assess freshwater condition through biomonitoring ^4–6^ and understand system stability and robustness to biodiversity loss and environmental stressors ^7–9^. Reproducible and scalable approaches for monitoring freshwater systems have never been more in demand than they are today.

Freshwater biomonitoring methods have evolved alongside the intensifying biodiversity declines, as demands grow for faster generation of mass data production (i.e. “big data”) ^10–12^. Typically, benthic macroinvertebrates are targeted for conducting freshwater health assessments, due to their taxonomic diversity, localized habitat occupancy and taxa-specific responses to a range of environmental gradients ^6, 12–15^. Across North America, reference sites are used for evaluations across watersheds, to account for variability in macroinvertebrate assemblages across ecoregions ^13, 16^. Region-specific tolerance values can then be generated via the Hilsenhoff Biotic Index (HBI), which provides a single tolerance value based on the average benthic arthropod community tolerance values to organic pollution (0 for very intolerant to 10 for highly tolerant) ^14, 17, 18^.

More recently, freshwater riverine microalgae, also referred to as diatoms, are also being used as bioindicators of rivers and streams, because of their strong response to environmental changes ^19–21^. Although microscopic morphological identification is currently the method of choice for diatom biomonitoring, high-throughput DNA metabarcoding of environmental samples has facilitated scaling up, primarily because of the ability of this method to bypass time-consuming morphology-based identifications ^22–25^. The combination of newly optimized and species-inclusive sample collection techniques (i.e. benthic kick-net ^25^) and reduced time taken to identify taxa ^22^ highlights the applicability of DNA metabarcoding as the ‘catch all’ approach for understanding freshwater condition. However, the ecological value of this ‘catch all’ method has not been investigated in real-world biomonitoring analyses.

DNA metabarcoding overcomes biomonitoring bottlenecks and enhances the amount of species-level diversity detected from environmental samples ^10, 11, 26–28^. The field is now in a position to move beyond simple biodiversity inventory measures such as richness, beta diversity, and community composition to associate species detections with known biological or ecological traits ^29–31^. One way to integrate and visualize this data, is through network analysis, a systems-level approach useful for integrating many layers of data. For example, metabarcoding data can be used to identify known trophic interactions, these interactions can then be used to build directed networks, food webs, to examine trophic relationships, identify keystone species, and clusters of potentially interacting species that can then be associated with their ecological traits such as tolerance to water pollution ^9, 12, 32–36^. Although trophic analysis is widely utilized in, for example, pollinator-plant, predator-prey systems; and network analysis has been widely utilized in the analysis of microbiome data ^37^ these approaches, are under-utilized in biomonitoring, particularly in river systems ^9^. Using species interactions to build trophic networks-can facilitate freshwater health assessments by visualizing the overall structural and functional relationships within a system ^38, 39^. More general network properties corresponding with ecosystem resilience and stability, such as connectedness and modularity, may also function like an early warning system for system collapse ^32, 40–44^.

Considering the role of multiple taxonomic groups (i.e. macroinvertebrates and diatoms) simultaneously as bioindicators can broaden the impact of freshwater assessments ^45, 46^. Multi-taxa approaches provide a more holistic representation of freshwater ecosystem health through the combination of taxonomic diversity, different species’ environmental tolerances and network properties (i.e. connectedness, modularity) from more than one traditional bioindicator kingdom ^45–48^. This ultimately enables the detection of keystone taxa that have a very large effect on their environment without which the community would be very different or not exist ^49^. Despite the evidence that multi-taxa biomonitoring approaches should be adopted, this is rarely the case due to logistical, time and cost restrictions involved with collecting representative samples for each taxonomic group ^46, 50^. There remains a lack of integration between DNA-based sample collection techniques for biomonitoring of macroinvertebrates and diatoms ^25^. The lengthy and multi-step nature of field sampling techniques for multiple taxa, can be overall detrimental to the amount of freshwater data collection ^50^ and is particularly incompatible with community-based monitoring (CBM), which is fast becoming a driving force for freshwater health data generation ^51^.

The objective of this study is to leverage trans-kingdom metabarcoding data generated from the same benthic kick-net samples to identify species associated with site condition in relation to known site condition and bioindicator taxa. Specifically, we: 1) use the cytochrome c oxidase subunit I (COI) (macroinvertebrate mitochondrial DNA marker) and ribulose bisphosphate large subunit (rbcL) (diatom chloroplast DNA marker) for the metabarcoding of benthic kick-net samples to generate biodiversity metrics (richness, effective number of exact sequence variants (ESVs), beta diversity) to assess subtly varying site condition (fair/good), 2) use multi-marker metabarcodes to identify site condition bioindicators for comparison with known stress tolerance, and 3) conduct an exploratory analysis of known trophic interactions to further assess the structure and stability of trophic networks across site conditions. We expect to determine unique bioindicators and keystone taxa, in addition to the well-known groups of bioindicator taxa because metabarcoding results are expected to both reflect and complement traditional sampling methods. We also predicted that ‘good’ quality sites would be more complex networks reflecting their ability to support a diverse array of taxa and functions.

## Results

A total of 3.2 million COI and 3.9 million rbcL sequence reads were generated for this study (Supplementary Tables 2 and 3). Following bioinformatic processing of raw reads, removing rare clusters, noise, chimeras, and pseudogenes a total of 4,026 COI and 1,573 rbcL ESVs (1,304,473 and 574,866 reads, respectively) were retained.

Rarefaction curves indicate that the sequencing depth was sufficient to capture the ESV diversity for both diatoms and macroinvertebrates across all four sites (Supplementary Fig. 1). After the COI and rbcL datasets were rarefied and normalized to the 15^th^ percentile of library sizes and merged, 45,937 reads in 2,933 ESVs were retained for further ESV level analyses.

### Diversity Analyses

At the order level, ‘fair’ sites show higher richness than ‘good’ sites for both diatoms and macroinvertebrates (Supplementary Fig. 3 & Supplementary Fig. 4). Diatoms from 12-17 orders were detected from samples from ‘fair’ sites and 9-16 orders from ‘good’ sites. Orders Naviculales, Cymbellales, Fragiliariales and Thalassiosirales were most prevalent across both ‘good’ and ‘fair’ sites. Genera within Thalassiosirales, Thalassiophysales and Bacillariales are known tolerant taxa ^52–54^ and had higher read abundance in ‘fair’ versus ‘good’ sites (Supplementary Fig. 3). We detected macroinvertebrates from 5 phyla: Platyhelminthes (flat worms), Nematoda (roundworms), Mollusca (molluscs), Arthropoda, and Annelida. Macroinvertebrates from 30-51 orders were detected from samples from ‘fair’ sites and 12-29 orders from ‘good’ sites. Traditional indicators of poorer water quality in river systems ^16^, including Haplotaxida, Gastropoda and Diptera and Odonata had higher read abundance in ‘fair’ sites. Although we appreciate that read abundance does not necessarily reflect organismal abundance in the environment, due to known issues with primer-bias and differential recovery of taxa with different body sizes, the relative abundance of these taxa across site conditions does correlate with what we would expect based on known species tolerances to pollution ^55, 56^.

The higher richness of ESVs compared to effective number of ESVs shows that the diversity across both site conditions is driven largely by many rare ESVs. The diversity detected in ‘fair’ sites was about twice as high in ‘good’ sites when measured using richness (macroinvertebrates: t-test, p.adj = 0.0039; diatoms: t-test, p.adj = 0.0240) but the effective number of ESVs were not found to be significantly different among site conditions (macroinvertebrates: t-test, p.adj = 0.92; diatoms: t-test. p.adj = 0.39; Fig. 1). Within each site condition diversity was similar between macroinvertebrates and diatoms (t-test, p.adj > 0.05; Fig. 1).

**Figure 1.**
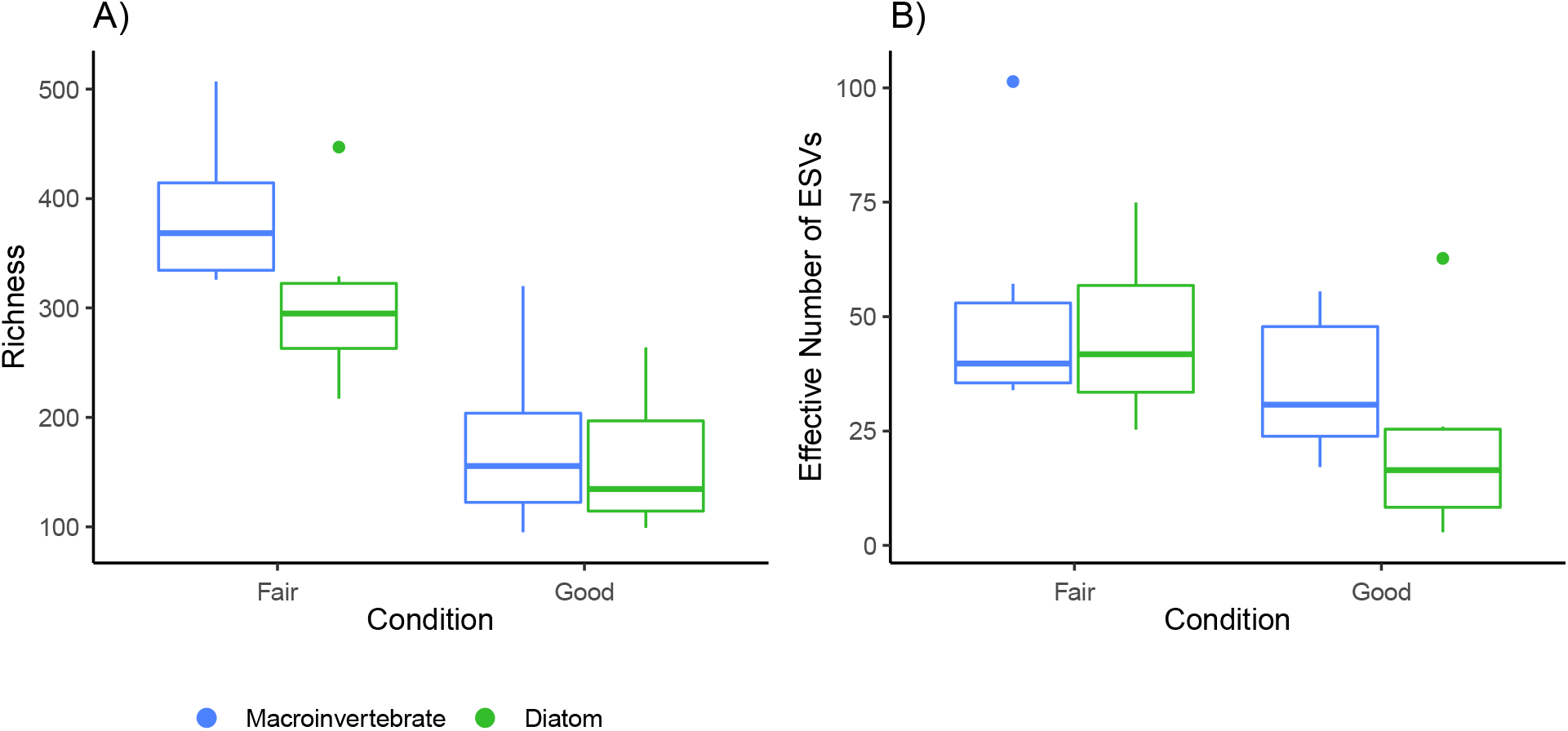
Observed richness is driven by a large number of rare ESVs. A) richness and B) The effective number of ESVs is shown for each condition and taxonomic group.

NMDS plots based on binary Bray-Curtis dissimilarities of ESVs across sites show good separation among fair and good sites using either diatoms or macroinvertebrates (diatoms stress = 0.04, linear R^2^ = 0.99; macroinvertebrates stress = 0.06, linear R^2^ = 0.98; Fig. 2). Diatoms and macroinvertebrates are both correlated with dissolved oxygen, turbidity, and pressure (mmHg). Additionally, macroinvertebrates are also correlated with pH and temperature. PERmutational ANalysis Of Variance (PERMANOVA) for diatoms showed that habitat status (good or fair) explained 22% of the variation in beta diversity (p-value = 0.001), whereas sampling site explains 25% of the variation (p-value = 0.004; Supplementary Table 4). For macroinvertebrates, habitat status explained 19% of the variation in beta diversity (p-value = 0.001), whereas site explained 35% of the variation (p-value = 0.002; Supplementary Table 4). The PERMANOVA reflects a combination of both dispersion (diatom site and status; macroinvertebrate site) and location effects as shown in the ordinations. Within each site condition (fair or good), dissimilarities were lower in fair sites and higher in good sites for diatoms but had a similar overlapping distribution for macroinvertebrates (Supplementary Fig. 5).

**Figure 2.**
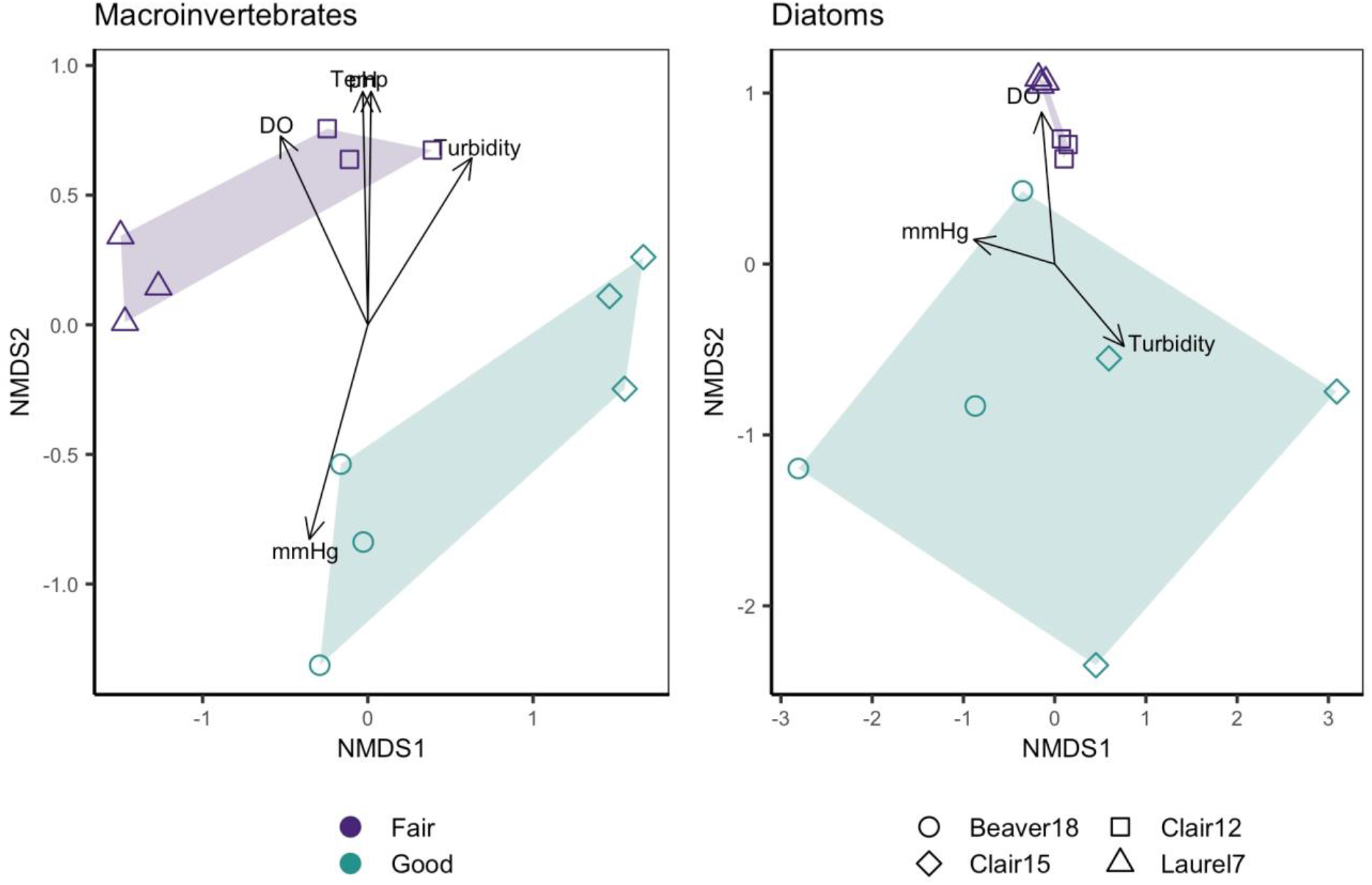
Beta diversity was greater within ‘good’ sites, especially for diatoms. Beta diversity within ‘fair’ sites, tended to be lower, especially for diatoms. Sample replicates cluster by site, and sites cluster by site status. Environmental variables that correlate with beta diversity patterns are shown if they have a p-value < 0.05. Based on binary Bray Curtis dissimilarities from libraries where read counts were rarefied to the 15^th^ percentile. Abbreviations: dissolved oxygen (DO); pressure (mmHg), temperature (Temp).

### Bioindicators

Results of indicator species analyses based on a rarefied read count matrix detected 29 site condition indicator species (Table 1). We recovered 28 fair condition indicators and one good condition indicator. The site condition indicators detected using the indicator value method (IndVal) and the point biserial correlation coefficient (*r*) were largely similar. The main differences between these statistics lie in their interpretation (Supplementary Fig. 6). The A and B components of the IndVal method indicate the predictive value and sensitivity/fidelity of the indicator ^57^. In this study, most of our site condition indicators have very high predictive value (close to 1) but only moderate sensitivity/fidelity. The point biserial correlation coefficient represents the ecological preference of species for a particular site condition and can range from -1 to +1, reflecting negative to positive correlations, and the closer to the absolute value of 1, the stronger the correlation ^58^. The advantage of including this measure, is that this method can detect both positive and negative correlations. This method recovered many of the same species as the indicator value method, all positively correlated with site condition. We also populated Table 1 with Biological Condition Gradient (BCG) scores from the Diatoms of North America Database (NADED; https://diatoms.org/) ^25^ and HBI scores for macroinvertebrates ^16^.

**Table 1.**
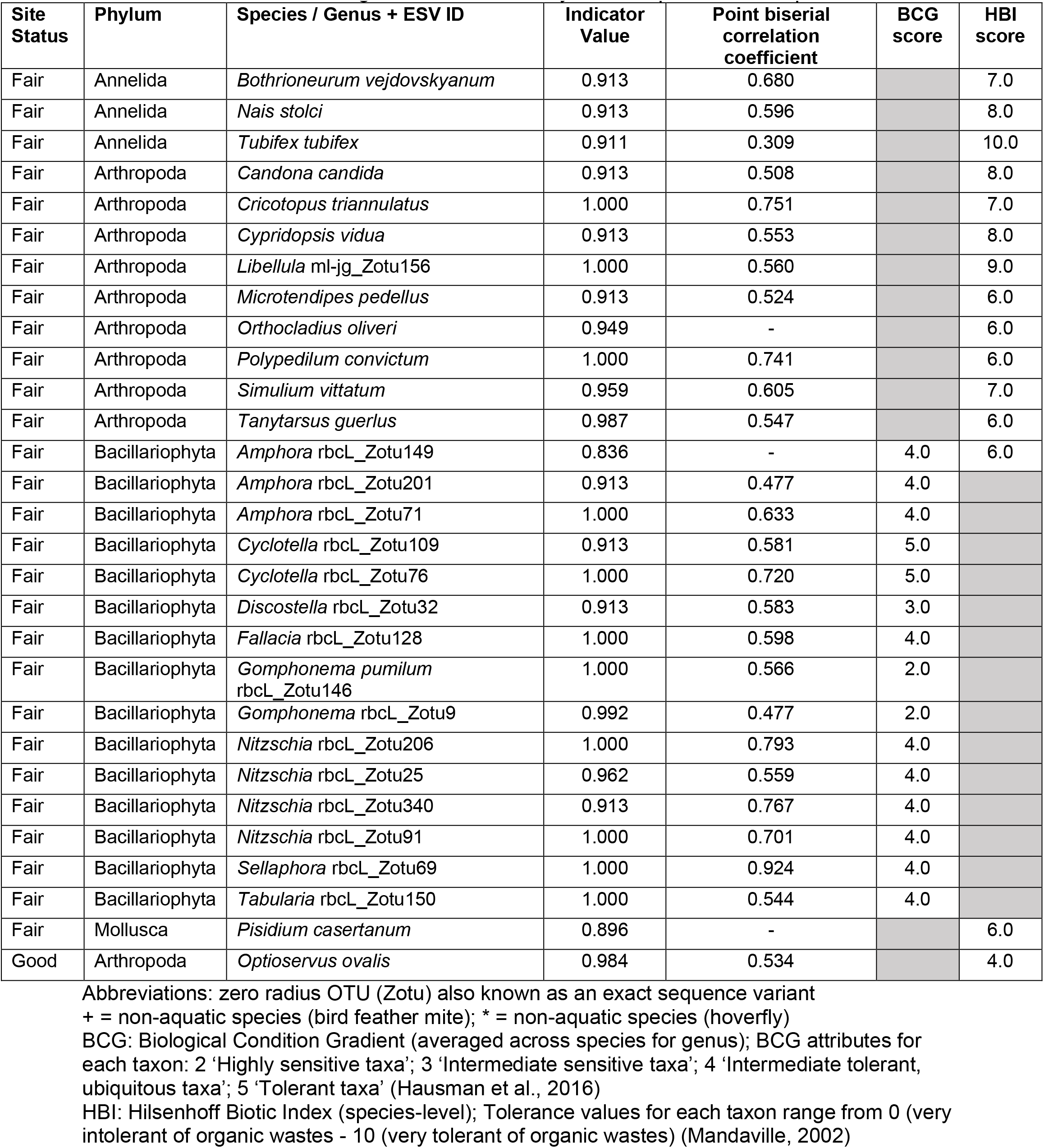
Diatom (n=15) and macroinvertebrate (n=17) habitat status indicator taxa. Based on rarefied data using ESV richness. Only shows species with a p-value < 0.05.

### Trophic interactions

An exploratory analysis using food webs for each site based on the automated retrieval of resource-consumer interactions was conducted (Fig. 3). GloBI annotation of resource to consumer interactions was possible for 71% (548/777) of our target taxa at the species and genus ranks. After filtering out interactions with off-target taxa (E.g., bacteria, fungi, plants, vertebrates), common names and insufficiently identified taxa (E.g., Chironomid, Lumbriculiid, Oligochaeta), and taxonomically unidentified substrates (E.g., CPOM - coarse particulate organic matter, detritus) target taxa representation was reduced to 34% (266/777). After filtering out off-target interactions, 22% (171/777) of our original target taxa were left represented in our interaction list. For each site, this means that 25.8 - 32.3% of the original target taxa were represented in each network. These trophic networks represent the current state of interaction annotations between diatoms and macroinvertebrates in GloBi and were used to visualize the trophic structure within each site and measure the network properties that would allow us to learn more about the stability of each site. Food webs generated from ‘fair’ habitat status sites tend to have more nodes (taxa), links (resource to consumer interactions), greater trophic height (longer food chains), and more clusters (Table 2). Food webs generated from ‘good’ habitat status sites, however, had slightly higher directed connectance (links/species^2^) and modularity (strength of divisions of a network into clusters). Similar to other described small-world type networks, our networks are highly clustered with relatively short path lengths ^59^. This means that most of the nodes in the network are not connected to each other, but the ones that are connected likely have neighbours that are also connected to each other, i.e., form clusters. Clair15 classified as a ‘good’ site had the smallest and sparsest food web, with lower numbers of nodes and trophic links. The Clair12 site classified as ‘fair’ had the greatest number of trophic links and trophic height. Fair condition sites tended to have more macroinvertebrate predators that are also tolerant to organic wastes (Table 3).

**Figure 3.**
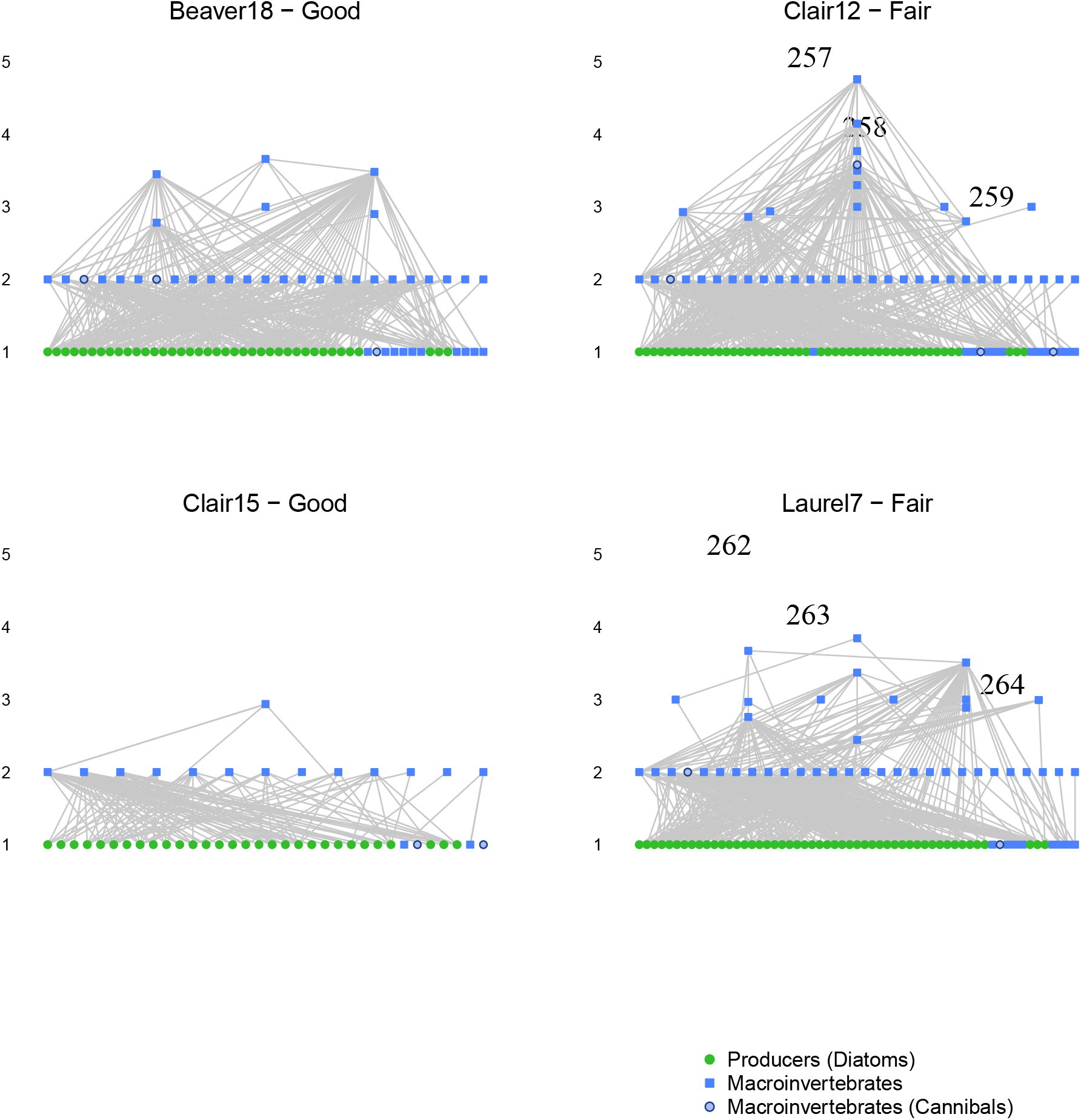
Food webs from ‘fair’ sites had higher trophic height and more macroinvertebrate predators. Vertical food webs with the lowest trophic level, producers (diatoms), at the bottom. Estimated trophic position for each node in the food web was determined using chain averaged trophic level (trophic height).

**Table 2.** ‘Good’ sites tended to have higher directed connectance and modularity whereas ‘fair’ sites tended to have a greater number of nodes, trophic links, and trophic height.

| Sites | Status | # Nodes | # Trophic links | Linkage density | Directed connectance | Characteristic path length | Trophic height | Modularity | Clusters* (total clusters) |
| --- | --- | --- | --- | --- | --- | --- | --- | --- | --- |
| Beaver 18 | Good | 83 | 316 | 3.8 | 0.05 | 2.4 | 3.7 | 0.08 | 5 (8) |
| Clair15 | Good | 45 | 106 | 2.4 | 0.05 | 2.2 | 2.9 | 0.16 | 4 (16) |
| Clair12 | Fair | 108 | 402 | 2.7 | 0.03 | 2.3 | 4.8 | 0.06 | 7 (10) |
| Laurel7 | Fair | 105 | 390 | 3.7 | 0.04 | 3 | 4.4 | 0.06 | 7 (15) |

\*Number of clusters with more than one taxon (total clusters including isolated taxa)

**Table 3.** More top predators that are tolerant to organic wastes are found in fair sites compared to good condition sites. Taxa with a trophic height greater than 3 are listed below, except for Clair 15 where the top macroinvertebrate predator is listed.

| Site Condition | Site | Predators (Trophic Height) | Functional Feeding Group<sup>1</sup> | Tolerance<sup>2,3</sup> |
| --- | --- | --- | --- | --- |
| Good | Beaver 18 | *Oecetis* (3.00)<br>*Hydropsyche* (3.48)<br>*Conchapelopia* (3.66) | PR<br>CG/CF/PR<br>PR | 5<br>1-6<br>6 |
| Good | Clair 15 | *Dicranota* (2.9375) | PR | 3 |
| Fair | Clair 12 | *Demicryptochironomus* (3.00)<br>*Parachironomus* (3.30)<br>*Procladius* (3.58)<br>*Hydropsyche* (3.59)<br>*Conchapelopia* (3.77)<br>*Enallagma* (4.18)<br>*Libellula* (4.80) | CG/PR<br>PR/CG/PA<br>PR/CG<br>CG/CF/PR<br>PR<br>PR<br>PR | 8<br>10<br>9<br>1-6<br>6<br>8<br>9<sup>3</sup> |
| Fair | Laurel 7 | *Conocephalus* (3.00),<br>*Oecetis* (3.00),<br>*Orconectes* (3.00),<br>*Carabus* (3.33),<br>*Libellula* (3.41),<br>*Hydropsyche* (3.51),<br>*Conchapelopia* (3.67),<br>*Argiope* (4.38),<br>*Graphoderus* (4.41) | PR<br>PR<br>CG/PR<br>PR<br>PR<br>CG/CF/PR<br>PR<br>PR<br>PR | -<br>5<br>6<br>-<br>9<br>1-6<br>6<br>-<br>5 |

<sup>1</sup> Functional feeding groups are from the EPA freshwater traits database or GloBI
<sup>2</sup> Ranges for tolerance values are from Mandaville (2002). Values range from 0 (intolerant to organic waste) to 10 (very tolerant to organic waste)
<sup>3</sup> For *Libellula* the tolerance for Libellulidae is given; ‘-’ indicates a tolerance value is unavailable; For *Orconectes* the tolerance for Cambaridae given

Directed graphs of resource-consumer interactions were also generated for each site (Fig. 4). This was done to identify clusters of potentially interacting taxa and to identify potential keystone taxa. Vertices (taxa) belonging to a cluster are encircled by a black line and the taxonomic composition of these clusters are detailed in Supplementary Tables 6-9. 4-5 clusters with more than one taxon were identified in ‘good’ sites and 7 clusters were identified from each ‘fair’ site (Table 2). Macroinvertebrate and diatom keystone genera for both ‘good’ and ‘fair’ sites were identified using two centrality measures: degree and hub scores and the top 3 scoring taxa from each site are summarized in Table 4. Degree and hub scores for all taxa are shown in Supplementary Tables 5-8. Generally, the distribution of degree and hub scores did not differ among sites assessed as ‘good’ or ‘fair’ (Supplementary Tables 5-8). While there were a few outlier invertebrates with a particularly high degree with many links to other taxa, diatoms in general tended to have higher hub scores than macroinvertebrates (Fig. 5). This reflects the large number of links from diatoms to a variety of macroinvertebrate consumers that themselves tend to feed on a variety of diatom species. Using the network terminology described by Kleinburg ^60^, this makes diatoms ‘hubs’ and diatom-consumers ‘authorities’. Diatoms play a key ecological role in linking macroinvertebrates together in trophic networks and this is reflected by the Kleinburg hub scores. Overall network modularity was assessed as low (0.1-0.15) to medium (0.15-0.2), being slightly higher for good sites (Table 2). In this study, we detected a greater number of clusters (containing more than one taxon) from networks with lower modularity. Though we detected more clusters, the strength of overall graph modularity was relatively weak, i.e. differential density of links within and between clusters not as stark.

**Figure 4.**
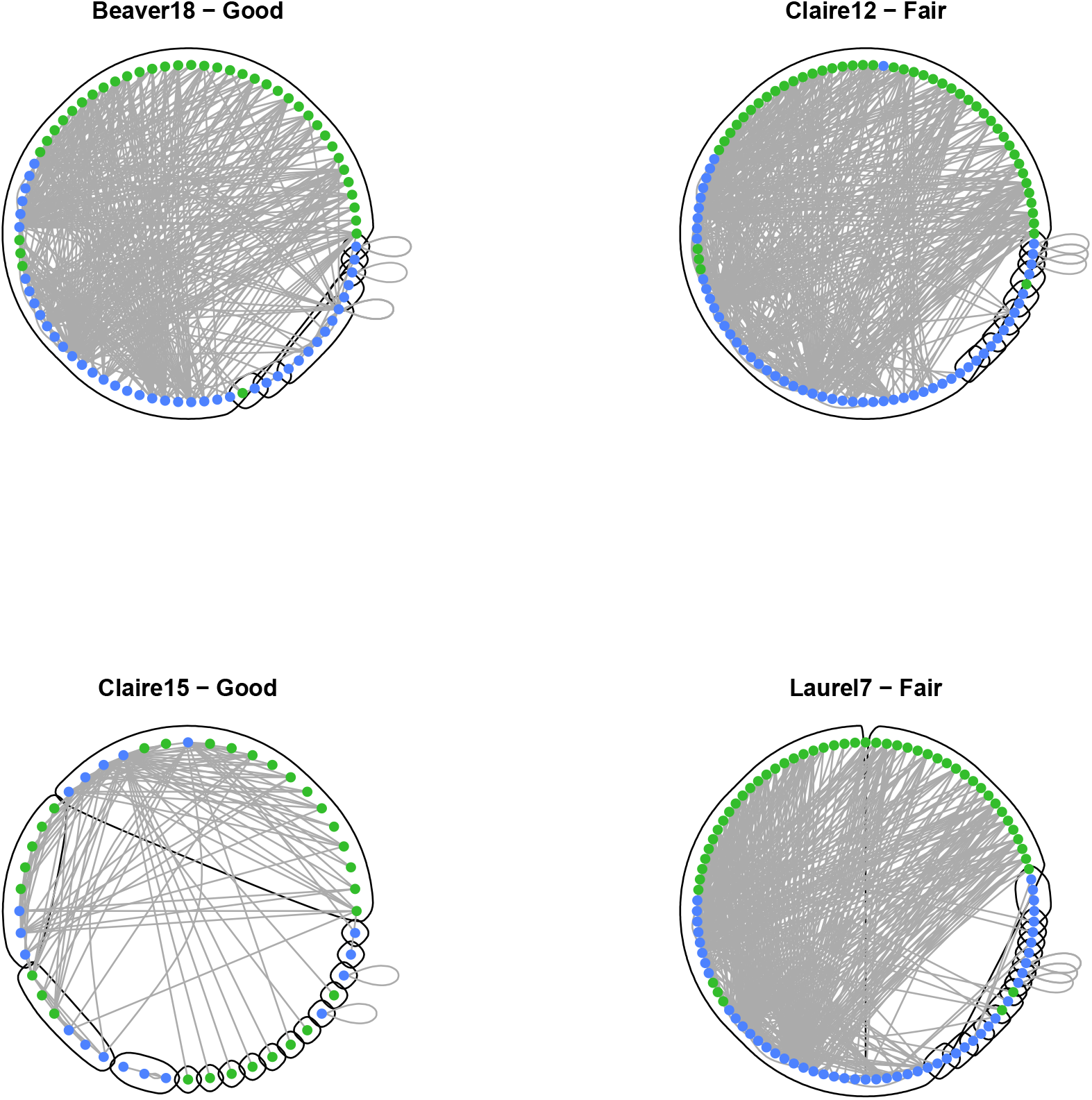
Freshwater benthic food webs show low to medium levels of modularity. Though these are directed graphs with interactions that point from resource to consumer, arrow heads were removed from the plot to improve readability. Trophic links (edges) are shown in grey. The small grey loops indicate cannibals, taxa known to consume members of the same taxon. Nodes (taxa) were arranged in a circle and colored according to trophic position (green-producers (diatoms); blue- macroinvertebrates). Clusters of taxa, as well as isolated taxa, are circled in black.

**Figure 5.**
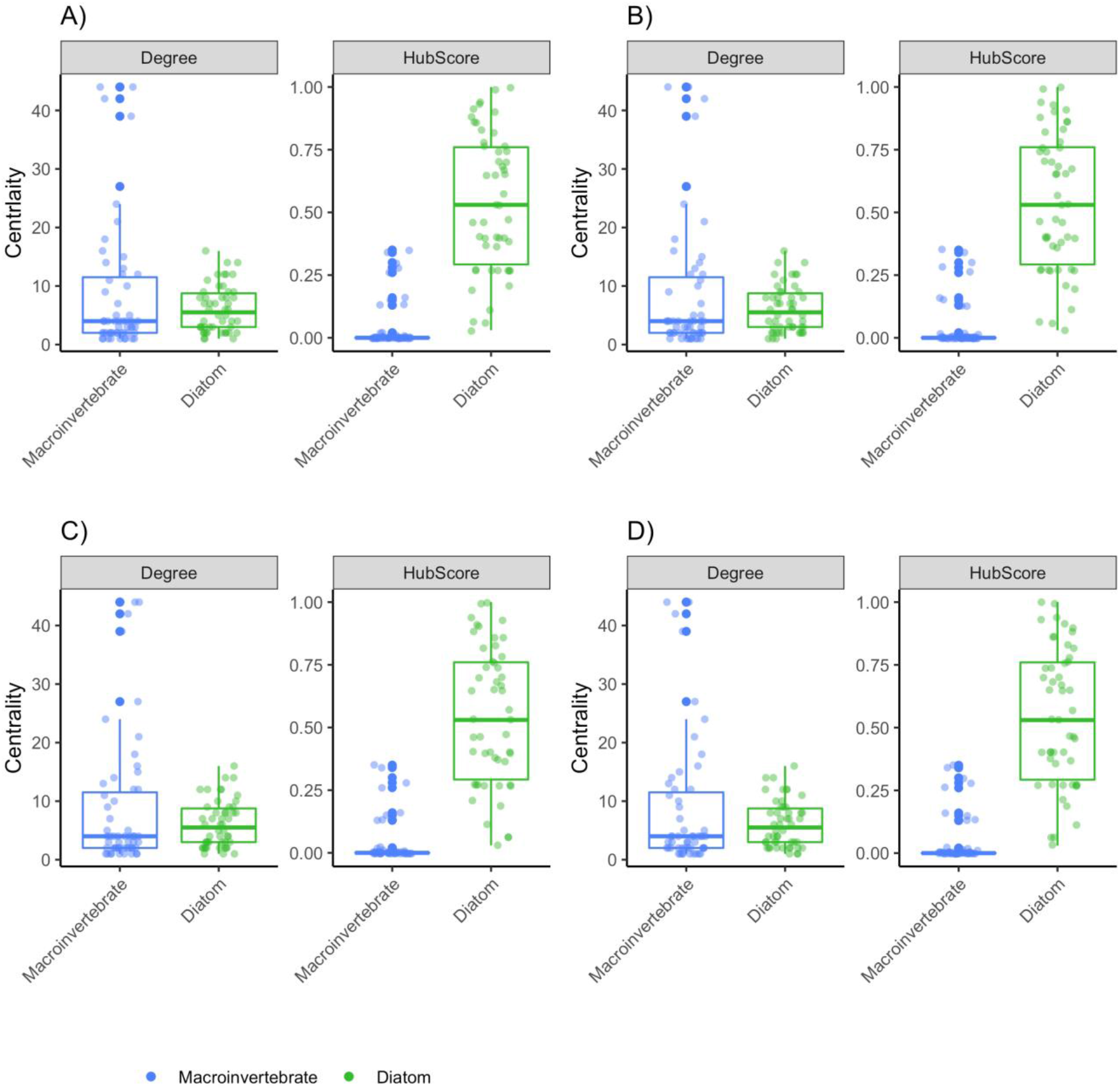
Kleinburg hub scores reflect the importance of diatoms in linking macroinvertebrate consumers together in trophic networks. Two measures used to detect potential keystone taxa are shown: 1) degree centrality and Kleinberg’s hub centrality scores. Degree shows the number of connections into and out of each taxon node. Hub scores reflect the number of connections out of each taxon node (hubs), from resource to consumer, weighted more havily when the consumers are themselves linked to by many other hubs. Sites are shown as follows: A) Beaver18 (good), B) Clair15 (good), C) Clair12 (fair), and D) Laurel7 (fair).

**Table 4.** Diatom (n=6) and macroinvertebrate (n=5) putative keystone taxa.

| Phylum | Genus* | Functional feeding guild | BCG Score | HBI score |
| --- | --- | --- | --- | --- |
| Bacillariophyta | *Navicula* |  | 4.0 |  |
| Bacillariophyta | *Fragilaria* |  | 3.0 |  |
| Bacillariophyta | *Rhoicosphenia* |  | 3.0 |  |
| Bacillariophyta | *Amphora* |  | 4.0 |  |
| Bacillariophyta | *Diatoma* |  | 2.0 |  |
| Bacillariophyta | *Gomphonema* |  | 2.0 |  |
| Arthropoda | *Cricotopus* (Chironomidae) | Collector-Gatherer |  | 7.0 |
| Arthropoda | *Hydropsyche* (Trichoptera) | Collector-Filterer |  | 4.0 |
| Arthropoda | *Polypedilum* (Chironomidae) | Shredder |  | 6.0 |
| Arthropoda | *Baetis* (Ephemeroptera) | Collector-Gatherer |  | 6.0 |
| Platyhelminthes | *Dugesia* | N/A |  | 6.0 |

\*Keystone taxa selected from top three degree and top three hub scores from each site
N/A: Not applicable
BCG: Biological Condition Gradient (averaged across species for genus)
HBI: Hilsenhoff Biotic Index (genus-level)

## Discussion

This study presents a trans-kingdom assessment of the parameters associated with a subtle variation in habitat status (fair/good). Multi-marker metabarcoding data detected both macroinvertebrates and diatoms from kick-net samples. We used this data to examine the biodiversity and network parameters associated with fair and good sites.

Macroinvertebrate bioindicator taxa are targeted globally as a means of identifying water quality status through assessment of assemblages in relation to environmental metadata ^13^. However, it can be difficult to understand aquatic health in relation to macroinvertebrate assemblages when evaluating ‘test’ sites which do not have corresponding reference sites and only have low-resolution regional tolerance values (e.g. to family and not species level) ^48^. This can be especially challenging for DNA metabarcoding-derived bioindicator species data, which requires tandem morphological-based studies for abundance assessments to be made^33^.

In our study, we utilized an integrated approach to rapidly identify site-specific bioindicator species. Through a combination of DNA-derived taxonomic assignments and indicator value/correlation analyses, we were able to determine site condition bioindicators without needing to identify species via morphology, or limit analyses to only EPT taxa. Biological Control Gradient (BCG) scores for the 15 diatom bioindicator species identified were between a 3 and 5, with the *Gomphonema* species being the only exception (BCG score of 2). A score between 3 and 5 is representative of an impaired system and reflects the point where diatom assemblages change due to increased human activity ^61^. *Gomphonema* as a genus which tends to be located within unimpaired systems ^62^, however, as this information is based on the ecoregion of California as opposed to eastern Canada ^63^, this may indicate that *Gomphonema* are more tolerant to poorer water quality in southern Ontario. Similarly, the Hilsenhoff Biotic Index (HBI) scores for identified ‘fair’ macroinvertebrate bioindicators ranged from 6 to 10, which falls within the ‘Fairly Poor’ to ‘Very Poor’ water quality categories ^16^. The one ‘good’ macroinvertebrate bioindicator species identified scored a HBI value of 4.0, whose presence in a system translates to ‘Very Good’ water quality status ^16^. Considering that the HBI was developed to detect organic pollution in aquatic systems through species weighting via relative abundance ^16, 17^, our site condition indicator species have been determined using rarefied read counts from metabarcoding data, without the need to quantify species abundance, and yet is still reflective of this index.

Despite the subtle habitat quality difference between the two site types, we have shown that it is still possible to identify site condition indicator species, especially for diatom taxa, which are lacking BCG metrics for Canadian systems ^61^. The habitat quality class used to assign our sites as ‘fair’ and ‘good’, are based on an amended HBI equation, which weighted each taxa present based on its tolerance value ^16, 25^. Unlike the HBI scores for macroinvertebrates, the BCG approach for assessing freshwater health and level of ecological impairment includes nutrient concentrations, other anthropogenic stressors, and possible confounding variables, and facilitates understanding of correlations between diatom assemblages and variables such as percentage of forest in watershed ^61^.

Beyond metrics of water quality, understanding the stability of aquatic ecosystem networks is important for predicting long-term resilience in the face of local and global environmental change scenarios ^64–67^. Generating networks of trophic interactions in freshwater systems can provide insight into ecosystem function, structure and robustness ^65, 68^. It has previously been shown that longer food webs are less stable and top predators more likely to go extinct ^69–71^. The trophic networks in our study show small-world characteristics, whereby most nodes are not neighbors of each other, but the neighbors of a node are likely to be neighbors of each other forming clusters ^59^. The short path lengths observed in our networks suggest that the effects of perturbations (e.g. species removal) would be distributed rapidly throughout the networks detected in our sites in a non-random fashion ^38, 59, 72^. Our networks also show relatively low modularity, with ‘good’ sites displaying marginally higher modularity. Higher network modularity is suggested to reflect higher stability, often through enhancing species persistence ^40, 73, 74^. By extension, lower network modularity may indicate that food webs in fair sites may be more susceptible to disturbance ^41^. In terms of effects of environmental stressors such as pollutants, more modular networks are likely to limit the propagation of both pollutants and their indirect effects through the food web ^75^. Despite higher richness in ‘fair’ sites, effective number of ESVs does not significantly differ among site conditions, indicating that rare taxa are more common in these ‘fair’ sites. Analyzing trait data for top predators at each site also indicates that ‘fair’ sites have more predators that also happen to be more resistant to poor water quality. Additionally, beta diversity, directed connectance, and modularity are all higher in ‘good’ sites, meaning these ‘good’ sites are expected to be more stable against persistent pollutant stress compared to ‘fair’ sites ^73, 75^.

In addition to determining stability through modularity, it is vital to determine presence of keystone taxa and trophic hubs, whose loss would likely cause cascading extinctions of many other species within freshwater food webs ^76^. Through targeting both diatoms and macroinvertebrates, we were able to determine keystone taxa from both producer and consumer trophic levels. Arthropod keystone taxa included genera from several traditional bioindicator groups (Ephemeroptera, Trichoptera and Chironomidae), of which perform a range of feeding strategies (e.g. collector-gatherer, collector-filterer and shredder). Despite diatoms often being excluded from network studies ^48^, the higher hub scores obtained for diatoms may reflect the importance of these producers as a food source for many different invertebrates ^77^. Several taxa such as *Amphora*, *Gomphonema*, *Crictopus*, and *Polypedilum* were identified as both keystone taxa and site condition bioindicators further reinforcing the ability of eDNA metabarcoding approaches to generate a robust picture of site condition and stability.

## Conclusion

There is an urgent need for more effective approaches to decipher biodiversity and ecosystem status as a consequence of environmental change especially due to global warming. Our study demonstrates that multi-taxa metabarcoding, is effective in identifying bioindicators of fine-scale freshwater condition and link these with their known tolerance to stressors. Unlike traditional methods, the use of multi-marker metabarcoding and indicator species analysis does not rely solely on the presence of EPT groups for making assessments of water quality, and enables a holistic measure of ecosystem health, even across previously identified subtle gradients of habitat quality status. While biodiversity analyses allowed us to distinguish site conditions based on alpha and beta diversity, correlation with environmental variables, as well as community composition, the addition of trophic network analyses also allowed us to identify clusters of taxa with known interactions, flag keystone taxa, and to assess ecosystem stability.

Trophic networks derived from eDNA data provide information on which key indicator interactions could signal a change in environmental conditions of a site, as opposed to only looking at presence/absence of traditional bioindicator taxa. Despite excluding leaf litter/detritus/fungi/microbes as resources for invertebrates in our food webs, we were still able to reconstruct highly connected systems and present trans-kingdom keystone taxa. In the same way that metabarcoding is considered a scalable approach to biomonitoring by automating the taxonomic assignment process; the annotation of trophic interactions also needs to be automated to be a scalable approach. The continued growth of online biotic interaction databases, from ecological studies, and text mining from the literature ^42, 78^, may one day help make the construction of global food webs a reality. Going forward, applying additional eDNA markers to target taxonomic groups such as fish, amphibians, and mammals, would greatly increase the level of trophic complexity in the networks and potentially identify additional bioindicator and keystone species, which may currently be overlooked in traditional biomonitoring strategies. Our work will set the stage for larger-scale studies involving sampling across a wide range of environmental gradients to further establish site condition bioindicator trends and potential influence of stressors on long-term ecosystem trophic networks.

## Methods

### Field Sampling

Samples were collected in November 2019 from Grand River tributaries across four study sites in Waterloo, Ontario (Supplementary Table 1; Supplementary Fig. 1). No specific permissions were required for sampling these sites because they are on public land and the field studies did not involve endangered or protected species. Status and location data were provided by Dougan & Associates based on a 2018 benthos biomonitoring project for the City of Waterloo (Supplementary Table 1). Clair15 and Clair12 are close in proximity, however Clair12 is directly downstream of several sewage outflows. The four selected sites were a subset of the sites from this project and were chosen based on accessibility and habitat quality. Hilsenhoff Biotic Index ranges (weighted by species) informed the habitat quality scale ^79^ which categorized sites into ‘Good’ (4.51-5.50) and ‘Fair’ (5.51-6.50).

Benthic kick-net samples were collected in triplicate within riffles, following the Canadian Aquatic Biomonitoring Network [CABIN] protocol ^80^, as previously described in ^25^. All samples were collected in 1L sample jars and placed in a cooler to transport back to the lab. Upon arrival at the lab, samples (n = 12) were preserved using >99% ethanol and stored in a -20°C freezer until processing.

### DNA Extraction

DNA from all samples was extracted following the methods previously detailed in ^25^. Briefly, samples were homogenized using blenders decontaminated with ELIMINase1 (Decon Labs, Pennsylvania, USA), rinsed with deionized water, and treated with UV light for 30 minutes. Homogenate was transferred to 50 mL Falcon tubes, one tube was centrifuged at 2400 rpm for two minutes. Supernatant was removed and pellets were dried at 70 °C. Approximately 300 mg dried tissue was used with the DNeasy Power Soil kit (Qiagen, CA) following the manufacturer’s protocol. Final elution was in 50 μL of buffer C6 (TE). Negative controls with no tissue were included with each batch of extractions. All negative controls failed to amplify and were not sequenced.

### DNA Amplification, Library Preparation and Sequencing Diatom rbcL

DNA amplification of samples for generation of diatom sequences is detailed in ^25^. Briefly, we targeted a 312 bp region of the chloroplast ribulose bisphosphate carboxylase large chain (rbcL) gene using five diatom specific primers: forward primers Diat_rbcL_708F_1, Diat_rbcL_708F_2 and Diat_rbcL_708F_3 combined in an equimolar mix; reverse primers Diat_rbcL_R3_1 and Diat_rbcL_R3_2 were also combined ^23^. The PCR cocktail was comprised of 17.5 μL HyPure^TM^ molecular biology grade water, 2.5 μL 10X reaction buffer (200 mM Tris-HCl, 500 mM KCl, pH 8.4), 1 μL MgCl_2_ (50 mM), 05. μL dNTPs mix (10 mM), 0.5 μL of both forward (10 mM) and reverse (10 mM) equimolar mixes, 0.5 μL Invitrogen’s Platinum Taq polymerase (5 U) and 2 μL of DNA for a final reaction volume totaled 25 μL. The PCR protocol was as follows: 35 cycles of denaturation at 95 °C for 45 seconds, annealing at 55 °C for 45 seconds and extension at 72 °C for 45 seconds. PCR amplification was also performed in two-steps: the first step used the taxon-specific primers listed above, with the second PCR used 2 μL of amplicons from the first PCR as template, with Illumina-adapter tailed taxon-specific primers. One negative PCR control was included with each PCR step, which both came back negative thus were not carried through to sequencing. All PCRs were completed in Eppendorf Mastercycler ep gradient S thermal cycler. Successful amplification was confirmed using 1.5% agarose gel electrophoresis before purifying second PCR amplicons with the MinElute Purification kit (Qiagen).

### Macroinvertebrate COI

Three fragments within the standard COI DNA barcode region were amplified with the following primer sets: (B/ArR5 [∼310 bp] called BR5, LCO1490/230_R [∼230 bp] called F230R, and mICOIintF/jgHCO2198 [∼313 bp] called ml-jg ^81–84^ using a two-step PCR amplification regime as described above, with the exception of the cycler conditions which were: initial denaturation of 95°C for 5min, 35 cycles of 94°C for 40s, 46°C for 1min and 72°C for 30s with a final extension of 72°C for 5min before holding at 10°C until PCRs were removed from the cycler.

Purified amplicons were quantified using a QuantIT PicoGreen daDNA assay kit and all samples were then normalised to 3 ng/µL, pooling the COI fragments for each sample before indexing with Ilumina Nextera adapters (FC-131-2001). Once indexed, samples were pooled into a single library and purified with AMpure magnetic beads. QuantIT PicoGreen dsDNA assay kit was once again used to quantify the library and Bioanalyzer was used to determine fragment length. rbcL and COI fragments were sequenced separately over two partial MiSeq runs. The purified libraries were diluted to 4 nM and sequenced according to manufacturers protocol, using a 10% PhiX spike-in before being sequenced using Illumina MiSeq with a V3 MiSeq sequencing kit (300 bp X 2; MS-102-2003).

### Bioinformatic Processing

Illumina MiSeq paired-end reads for both COI and rbcL were processed using the MetaWorks-1.3.1 pipeline ^85, 86^ available from https://github.com/terrimporter/MetaWorks. MetaWorks is an automated Snakemake ^87^ bioinformatic pipeline that runs in a conda ^88^ environment. SeqPrep v1.3.2 ^89^ was used to pair raw reads requiring a minimum Phred score of 20 in the overlap region to ensure 99% base-calling accuracy and a minimum of 25 bp overlap. CUTADAPT v2.6 was used to trim primers from sequences, using a Phred score cutoff of 20 at the ends, leaving a minimum fragment length of at least 150 base pairs, no more than 3 N’s permitted ^90^. Global exact sequence variants (ESV) ^91^ were generated for the primer-trimmed reads. Reads were dereplicated using the ‘derep_fulllength’ command with the ‘sizein’ and ‘sizeout’ options of VSEARCH v2.14.1 ^92^. VSEARCH was also used to denoise the data using the unoise3 algorithm ^93^. These steps were taken to remove sequences with errors and rare reads (clusters with only one or two reads) ^94^. Putative chimeric sequences were removed using the ‘uchime3_denovo’ algorithm in VSEARCH. An ESV x sample table was created using the ‘search_exact’ method in VSEARCH.

Diatom rbcL ESVs were classified using the rbcL diatom reference set available from https://github.com/terrimporter/rbcLdiatomClassifier ^25, 95^. The reference sequence set is based on rbcL sequences from the Diat.barcode project ^96, 97^ and reformatted to train a naive Bayesian classifier to make rapid, accurate taxonomic assignments ^98^. This method makes assignments to the species rank and produces a statistical measure of confidence for each taxon up to the domain rank to help reduce false positive taxonomic assignments. Species level assignments used a 90% bootstrap support cutoff, no cutoff was needed at the genus rank, to expect at least 90% correct taxonomic assignments assuming the query sequences are represented in the reference database. Macroinvertebrate COI ESVs were classified using the COI Classifier v4 available from https://github.com/terrimporter/CO1Classifier/releases/tag/v4 ^99^, comprised of a curated reference sequence set mined from BOLD ^100^ and GenBank ^101^, and uses the RDP classifier v2.12 that uses a naive Bayesian algorithm ^98, 102^. We used a 0.70 bootstrap support cutoff at the species rank (90% correct), and no cutoff at the genus rank was needed, to expect 95% correct taxonomic assignments assuming the query sequences are represented in the reference database.

As we were using protein coding markers in this study, we also screened out obvious pseudogenes to try to reduce noise in the dataset and avoid inflating richness estimates ^103^. For rbcL, we removed putative pseudogenes using removal method 1: rbcL ESVs were translated into every possible reading frame, plus strand only, using ORFfinder v0.4.3, keeping the longest open reading frame (ORF). ORFs with shorter or longer outlier sequence lengths were removed as putative pseudogenes. For COI, putative pseudogenes were identified and removed in the METAWORKS pipeline using removal method 2 since a hidden Markov model (HMM) was available for this marker. COI ESVs were translated into ORFs as described above, and for each ESV, the longest ORF was retained. Amino acid ORFs were used for HMM profile analysis and ORFs with low outlier sequence bit scores were removed as putative pseudogenes.

### Data Analyses

RStudio v1.3.1093 was used with R v4.0.3 to analyze the data ^104, 105^. Plots were created with ggplot2 except for specialized plots where indicated ^106^. Sites were plotted using the ‘ggmap’ package with stamen maps (Map tiles by Stamen Design, under CC BY 3.0. Data by OpenStreetMap, under ODbL. Available from http://maps.stamen.com/#watercolor/12/37.7706/-122.3782 ) ^107^.To account for variable reads in each library, sample read number was normalized to the 15^th^ percentile library size using the ‘rrarefy’ function in the vegan package ^108, 109^. Rarefaction curves were plotted using a slightly modified ‘rarecurve’ function (Supplementary Fig. 2). Sequencing depth was found to be sufficient to capture amplicon diversity across samples, as each curve reached a plateau, even after rarefying to the 15^th^ percentile read depth. Unless otherwise stated, all further analyses are based on rarefied data.

We checked for differences in richness, the average number of ESVs per sample, site conditions (fair or good, 2 sites per condition, 3 replicates per site), and taxonomic groups (Arthropoda, Bacillariophyta). However, since richness is insensitive to species frequencies (rare species are weighted equally to common species), we also calculated the numbers equivalent, i.e., the effective number of equally-frequent species, using the exponential of Shannon entropy ^110^ in the vegetarian package ^111^, which is less sensitive to rare ‘species’. Our richness and effective number of ESVs were found to be normally distributed using a quantile-quantile plot using the ggqqplot function in R as well as the Shapiro-Wilk test using the shapiro.test function in R (p = 0.27). T-tests were used to compare sample means across site conditions and across taxa within site conditions. The Holm method was used to adjust for multiple comparisons. To illustrate the range of biodiversity detected using multi-marker metabarcoding, as well as among-sample variability, we plotted a heatmap to visualise the detection of diatom and macroinvertebrate orders (Supplementary Fig. 3 and Supplementary Fig. 4). We reviewed our taxa and removed any non-freshwater taxa from our indicator lists. This may due to taxonomic misidentification in the underlying reference database ^112^.

A non-metric multi-dimensional scaling (NMDS) analysis on binary Bray-Curtis (Sorensen) dissimilarity matrix was conducted using the vegan ‘metaMDS’ function ^113^. A scree plot was run using the ‘dimcheckMDS’ command from the goeveg package to determine the number of dimensions (k=2) to use with the vegan metaMDS function ^114^. Shephard’s curve and goodness of fit plots were created using the vegan ‘stressplot’ and ‘goodness’ functions. The vegan ‘vegdist’ command was used to build a binary Bray Curtis dissimilarity matrix. We checked for heterogeneous distribution of dissimilarities using the ‘betadisper’ function. We used the ‘adonis’ function to perform permutational multivariate analysis of variance (PERMANOVA). PERMANOVA ^115^ was performed to assess whether sites or site status had any significant interactions or explained any variation in beta diversity. Environmental variables (temperature, percentage dissolved oxygen, pressure, specific conductance, pH and turbidity) were fitted to the NMDS plot using the ‘envfit’ function in vegan with 999 permutations. Only significant variables (p<0.05) were plotted.

We used the ‘multipatt’ function from the indicspecies R package to: identify species that could be used as indicators of site quality using the Indicator Value method (IndVal); and to identify species correlated with environmental conditions at fair and poor sites using the point biserial correlation coefficient ^57, 58^. Both functions used the “.g” version to correct for unequal group sizes and we set duleg=TRUE to avoid considering site group combinations. We included taxa at the species rank where we could, otherwise we retained the genus level assignment and appended the ESV ID. Each test was run using a taxon x sample table containing rarefied read counts. We analyzed the diatom and macroinvertebrate species assemblages independently, to determine the strongest indicators within each taxonomic group.

For trophic analyses, we worked with taxa at the species and genus rank as described above except that we did not append ESV IDs to genera. The list of target taxa was manually reviewed and edited to account for insufficiently identified species assignments with ‘sp.’, ‘cf.’, or an alphanumeric code ^116^. We also summarized identifications at the variety level, containing ‘var.’, to the species rank. We obtained biotic interactions for each taxon from the Global Biotic Interactions (GloBI) database ^31^ using the ‘get_interactions_by_taxa’ function in the rglobi package in R ^117^. In our first search, we retrieved interactions for some of the species and genera in our target list. For species that were not detected, we collapsed the taxonomic assignment to the genus rank and conducted a second search. For all searches conducted at the genus rank, the retrieved interactions were pooled across the species within the genus. We filtered interactions to only keep ones where both the resource and consumer were detected in a site. We then filtered the interactions to only keep those that described the directed resource to consumer relationship (ex. “eatenBy”, “preyedUponBy”). With the cheddar package, we used our directed resource to consumer interactions to calculate several measures of trophic complexity including number of nodes (species), trophic links (interactions), linkage density (links/species), directed connectance (links/species^2^), characteristic path length (average of path lengths from each node to a basal species), as well as trophic height (chain averaged trophic length) ^118–120^. The food webs were visualized using the ‘PlotWebByLevel’ function with the chain averaged trophic level method.

For network analysis, we used the same resource to consumer relationships from above to build directed graphs using the ‘graph.data.frame’ function in the igraph package. We used the walktrap method to identify communities of potentially interacting (co-occurring) taxa, also referred to as modules, clusters, groups, or subgraphs in the literature. The ‘cluster_walktrap’ function identifies clusters via random walks, with the assumption that short random walks tend to stay in the same community, and edges within a cluster are denser than edges between clusters. For each site, we also assessed overall network modularity, the strength of divisions of a network into modules (clusters). We categorized modularity as follows: very low (< 0.1), low (0.1-0.15), medium (0.15-0.2), high (0.2-0.3), and very high (> 0.3) ^121^. We used two different measures to identify keystone taxa, that is, nodes with high centrality. First, we calculated degree by recording the number of edges (co-occurrences) into and out of a vertex (species). Second, we calculated Kleinberg’s hub centrality scores (hub scores) that takes into consideration the authority and hubbiness of a vertex ^60^. An authority refers to the number of links into a vertex and hubbiness refers to the number of links out to other vertices with high authority. Degree and hub scores were calculated for every taxon in the network and for each measure, the top 3 taxa from each site were retained as a potential keystone taxon. The network was circularized with vertices ordered by cluster membership.

## Acknowledgements

This study is funded by the Government of Canada through Genome Canada and Ontario Genomics. Teresita Porter was funded by the Government of Canada through the Genomics Research and Development Initiative (GRDI) Metagenomics-Based Ecosystem Biomonitoring Project (Ecobiomics).

## Author Contributions and Competing Interests

C.V.R and M.H. designed the study, C.V.R contributed to sampling, bioinformatic processing and statistical analyses, T.M.P created the bioinformatic pipeline and trained the classifiers, contributed to bioinformatic processing and performed data analysis, V.C.M. collected samples and conducted molecular analyses, M.T.G.W. conducted molecular analysis and sequenced the samples. All authors helped to write/edit the manuscript.

## Additional Information

The authors have declared that no competing interests exist.

## Data Availability

Raw sequences will be available from NCBI SRA on acceptance. The MetaWorks-1.3.1 is available from https://github.com/terrimporter/MetaWorks, the rbcLdiatomClassifier v1 and COIClassifier v4 we used are available on GitHub at <u>ttps://github.com/terrimporter/rbcLdiatomClassifier</u> and https://github.com/terrimporter/CO1Classifier. Scripts and files used to generate outputs can be found at https://github.com/terrimporter/RobinsonEtAl2021_MacroinvertDiatom.

