## Supplementary Information for "Multi-marker metabarcoding resolves subtle variations in freshwater condition: Bioindicators, ecological traits, and trophic interactions"

**Supplementary Table 1. Information on study sites, including GPS coordinates, site status based on the Hilsenhoff Biotic Index ranges (weighted by species) and environmental metadata collected in 2019.**

| Site | Latitude | Longitude | Site status | Temp °C | Hg (mmHg) | DO % | Sp Cond | pH | Turbidity FNU |
|---|---|---|---|---|---|---|---|---|---|
| Clair15 | 43.462901 | -80.58468 | Good | 5.6 | 731.6 | 81.9 | 383.6 | 8.29 | 27.63 |
| Beaver18 | 43.492001 | -80.60978 | Good | 4.3 | 737.7 | 83.5 | 578 | 7.99 | 2.19 |
| Laurel7 | 43.470727 | -80.55627 | Fair | 5.6 | 734.6 | 88.3 | 606 | 8.36 | 8.95 |
| Clair12 | 43.465409 | -80.57132 | Fair | 9.2 | 733.1 | 88.4 | 3978 | 8.54 | 13.60 |

Abbreviations: Lat = Latitude; Lon = Longitude; Temp = Temperature; Hg = Pressure; DO = Dissolved oxygen; Sp Cond = Specific conductance

**Supplementary Table 2. COI read and ESV statistics.**

| Step | BR5 | F230R | ml-jg | Total |
|---|---|---|---|---|
| **Raw paired-end reads** | - | - | - | 3,254,709 x 2 |
| **Paired reads** | - | - | - | 3,130,774 |
| **Primer trimmed reads** | 648,685 | 951,955 | 1,360,596 | |
| **ESVs (reads)** | 575 (131,576) | 923 (338,612) | 955 (259,419) | 4,026 (1,304,473) |

**Supplementary Table 3. rbcL read and ESV statistics.**

| **Step** | **rbcL** |
|---|---|
| Raw paired-end reads | 3,918,107 x 2 |
| Paired reads | 3,607,536 |
| Primer trimmed reads | 1,652,415 |
| ESVs (reads) | 1,573 (574,866) |

**Supplementary Table 4. Both diatoms and macroinvertebrates separate fair and good sites.** Significant beta dispersion was detected for status and sites for diatoms; and for site for macroinvertebrates. Summary of PERMANOVA results based on a Sorensen dissimilarity matrix of diatom and macroinvertebrate order level ESVs. P-values < 0.05 are in bold.

| **Taxon** | **Formula / Source of variation** | **Df** | **SS** | **MS** | **F** | **$R^2$** | **P** |
|---|---|---|---|---|---|---|---|
| **Diatoms** | diss ~ Status * Site | | | | | | |
| | Status | 1 | 0.810 | 0.810 | 3.312 | 0.219 | **0.001** |
| | Site | 2 | 0.939 | 0.470 | 1.920 | 0.253 | **0.004** |
| | Residuals | 8 | 1.957 | 0.245 | | 0.528 | |
| | Total | 11 | 3.707 | | | 1.000 | |
| **Macroinvertebrates** | diss ~ Status * Site | | | | | | |
| | Status | 1 | 0.797 | 0.797 | 3.302 | 0.190 | **0.001** |
| | Site | 2 | 1.465 | 0.732 | 3.036 | 0.349 | **0.001** |
| | Residuals | 8 | 1.930 | 0.241 | | 0.460 | |
| | Total | 11 | 4.192 | | | 1.000 | |

Abbreviations: Sums of Squares (SS); Mean Squares (MS); diss (binary Bray Curtis dissimilarities among samples); Site (Clair 12, Clair 15, Beaver 18 or Laurel 07); Status (Good or Fair)

**Supplementary Table 5. Beaver18 (good) clusters and keystone taxa (in bold).**

Members of the same group were given the same numeric designation.

| Category | Taxon | Group | Vertex Number | Degree | Hub Score |
|---|---|---|---|---|---|
| **Producer** | **Navicula**** | 1 | 1 | 15 | **0.9** |
| Producer | Navicula gregaria | 1 | 2 | 4 | 0.36 |
| Producer | Navicula lanceolata | 1 | 3 | 4 | 0.36 |
| Producer | Meridion circulare | 1 | 4 | 10 | 0.42 |
| Producer | Amphora pediculus | 1 | 5 | 5 | 0.4 |
| **Producer** | **Fragilaria**** | 1 | 6 | 17 | **0.93** |
| Producer | Caloneis | 1 | 7 | 8 | 0.56 |
| Producer | Gomphonema truncatum | 1 | 8 | 3 | 0.22 |
| Producer | Planothidium lanceolatum | 1 | 9 | 3 | 0.31 |
| Producer | Rhoicosphenia abbreviata | 1 | 10 | 7 | 0.46 |
| Producer | Epithemia | 1 | 11 | 9 | 0.55 |
| Producer | Nitzschia linearis | 1 | 12 | 1 | 0.05 |
| Producer | Cymbella | 1 | 13 | 13 | 0.84 |
| Producer | Surirella brebissonii | 1 | 14 | 1 | 0.1 |
| Producer | Nitzschia dissipata | 1 | 15 | 5 | 0.44 |
| Producer | Cocconeis placentula | 1 | 16 | 10 | 0.52 |
| Producer | Melosira varians | 1 | 17 | 5 | 0.38 |
| Producer | Meridion | 1 | 18 | 5 | 0.33 |
| Producer | Navicula tripunctata | 1 | 19 | 4 | 0.36 |
| Producer | Cocconeis | 1 | 20 | 10 | 0.55 |
| Producer | Navicula cryptocephala | 1 | 21 | 2 | 0.17 |
| Producer | Staurosira | 1 | 22 | 10 | 0.7 |
| Producer | Diploneis | 1 | 23 | 5 | 0.46 |
| Producer | Frustulia | 1 | 24 | 8 | 0.61 |
| Producer | Neidium | 1 | 25 | 4 | 0.37 |
| Producer | Gomphonema | 1 | 26 | 13 | 0.85 |
| Producer | Encyonema | 1 | 27 | 10 | 0.62 |
| Producer | Nitzschia | 1 | 28 | 8 | 0.64 |
| **Producer** | **Rhoicosphenia**** | 1 | 29 | 18 | **1** |
| Producer | Hannaea | 1 | 30 | 8 | 0.53 |
| Producer | Pinnularia | 1 | 31 | 7 | 0.59 |
| Producer | Achnanthidium | 1 | 32 | 4 | 0.37 |
| Producer | Psammothidium | 1 | 33 | 6 | 0.51 |
| Producer | Eunotia | 1 | 35 | 7 | 0.62 |
| Producer | Bacillaria | 1 | 36 | 4 | 0.37 |

| **Category** | **Taxon** | **Group** | **Vertex Number** | **Degree** | **Hub Score** |
|---|---|---|---|---|---|
| Invertebrate | Polypedilum | 1 | 37 | 25 | 0.1 |
| Invertebrate | Chironomus | 1 | 38 | 1 | 0.08 |
| Invertebrate | Tanytarsus | 1 | 39 | 12 | 0.31 |
| Invertebrate | Micropsectra | 1 | 41 | 6 | 0.05 |
| Invertebrate | Simulium | 1 | 43 | 18 | 0.36 |
| **Invertebrate** | **Cricotopus*** | 1 | 46 | **35** | 0.32 |
| Producer | Planothidium | 1 | 50 | 8 | 0.63 |
| Producer | Sellaphora | 1 | 51 | 4 | 0.37 |
| **Producer** | **Amphora**** | 1 | 52 | 15 | **0.9** |
| Invertebrate | Tipula | 1 | 53 | 10 | 0 |
| Invertebrate | Orthocladius | 1 | 54 | 11 | 0.27 |
| Invertebrate | Prosimulium | 1 | 55 | 9 | 0.03 |
| Invertebrate | Thienemanniella | 1 | 61 | 5 | 0.14 |
| Invertebrate | Baetis | 1 | 62 | 27 | 0.31 |
| Invertebrate | Nais | 1 | 63 | 2 | 0.27 |
| Invertebrate | Hyalella azteca | 1 | 64 | 5 | 0 |
| **Invertebrate** | **Hydropsyche*** | 1 | 65 | **47** | 0 |
| Invertebrate | Pisidium | 1 | 66 | 14 | 0 |
| Invertebrate | Pseudokiefferiella | 1 | 67 | 3 | 0 |
| Invertebrate | Diamesa | 1 | 68 | 10 | 0 |
| Invertebrate | Simulium vittatum | 1 | 69 | 5 | 0 |
| Invertebrate | Antocha saxicola | 1 | 70 | 1 | 0 |
| Invertebrate | Stenelmis crenata | 1 | 71 | 1 | 0 |
| Invertebrate | Lype | 1 | 72 | 21 | 0 |
| Invertebrate | Lepidostoma | 1 | 73 | 21 | 0 |
| Invertebrate | Prodiamesa | 1 | 74 | 5 | 0 |
| Invertebrate | Cloeon | 1 | 75 | 21 | 0 |
| Invertebrate | Lymnaea | 1 | 76 | 8 | 0 |
| Invertebrate | Calanus | 1 | 77 | 2 | 0 |
| Invertebrate | Oecetis | 1 | 79 | 2 | 0 |
| Producer | Gomphoneis | 2 | 34 | 3 | 0.17 |
| Invertebrate | Stempellinella | 2 | 78 | 1 | 0 |
| Invertebrate | Chrysops | 3 | 58 | 1 | 0 |
| Invertebrate | Conocephalus | 3 | 81 | 2 | 0 |
| Invertebrate | Chaetocladius | 4 | 42 | 1 | 0.05 |
| Invertebrate | Nanocladius | 4 | 48 | 1 | 0.05 |
| Invertebrate | Conchapelopia | 4 | 49 | 6 | 0.13 |
| Invertebrate | Rheotanytarsus | 4 | 56 | 5 | 0.36 |
| Invertebrate | Paratanytarsus | 4 | 57 | 2 | 0.05 |
| Invertebrate | Glyptotendipes | 4 | 59 | 2 | 0.09 |

| Category | Taxon | Group | Vertex Number | Degree | Hub Score |
|---|---|---|---|---|---|
| **Invertebrate** | **Enallagma*** | 4 | 60 | **28** | 0.14 |
| Invertebrate | Fannia | 5 | 47 | 1 | 0 |
| Invertebrate | Hydrotaea | 5 | 80 | 1 | 0 |
| Invertebrate | Drosophila | 6 | 40 | 2 | 0 |
| Invertebrate | Sperchon | 7 | 44 | 1 | 0.01 |
| Invertebrate | Stagmomantis | 8 | 45 | 4 | 0 |

* in the top 3 based on degree

** in the top 3 based on hub score

**Supplementary Table 6. Clair12 (fair) clusters and keystone taxa (in bold).**

Members of the same group were given the same numeric designation.

| Category | Taxon | Group | Vertex Number | Degree | Hub Score |
|---|---|---|---|---|---|
| Producer | Navicula gregaria | 1 | 1 | 8 | 0.62 |
| Producer | Navicula lanceolata | 1 | 2 | 6 | 0.52 |
| **Producer** | **Diatoma**** | 1 | 3 | 17 | **0.9** |
| Producer | Diatoma vulgaris | 1 | 4 | 1 | 0.11 |
| Producer | Amphora pediculus | 1 | 5 | 10 | 0.68 |
| **Producer** | **Fragilaria**** | 1 | 6 | 21 | **0.99** |
| Producer | Gomphonema truncatum | 1 | 7 | 4 | 0.35 |
| Producer | Rhoicosphenia abbreviata | 1 | 8 | 10 | 0.65 |
| Producer | Sellaphora seminulum | 1 | 9 | 1 | 0.11 |
| Producer | Amphora ovalis | 1 | 10 | 8 | 0.54 |
| Producer | Nitzschia linearis | 1 | 11 | 1 | 0.06 |
| Producer | Navicula cryptotenella | 1 | 12 | 1 | 0.11 |
| Producer | Cymbella | 1 | 13 | 10 | 0.72 |
| Producer | Nitzschia amphibia | 1 | 14 | 1 | 0.11 |
| Producer | Surirella brebissonii | 1 | 15 | 2 | 0.21 |
| Producer | Nitzschia dissipata | 1 | 16 | 8 | 0.65 |
| Producer | Cocconeis placentula | 1 | 17 | 17 | 0.88 |
| Producer | Melosira varians | 1 | 18 | 9 | 0.62 |
| Producer | Navicula tripunctata | 1 | 19 | 8 | 0.62 |
| Producer | Cymbella tumida | 1 | 20 | 1 | 0.06 |
| Producer | Navicula capitatoradiata | 1 | 21 | 1 | 0.11 |
| Producer | Cocconeis | 1 | 22 | 10 | 0.61 |
| Producer | Achnanthes | 1 | 23 | 16 | 0.85 |
| Producer | Navicula cryptocephala | 1 | 24 | 3 | 0.29 |
| Invertebrate | Macrocyclops albidus | 1 | 25 | 5 | 0.21 |
| Producer | Navicula | 1 | 26 | 14 | 0.89 |
| Producer | Diploneis | 1 | 27 | 3 | 0.34 |
| Producer | Frustulia | 1 | 28 | 8 | 0.55 |
| Producer | Neidium | 1 | 29 | 2 | 0.24 |
| Producer | Caloneis | 1 | 30 | 5 | 0.44 |
| Producer | Gomphonema | 1 | 31 | 13 | 0.86 |
| Producer | Encyonema | 1 | 32 | 6 | 0.47 |
| Producer | Nitzschia | 1 | 33 | 6 | 0.52 |
| **Producer** | **Rhoicosphenia**** | 1 | 34 | 19 | **1** |

| Category | Taxon | Group | Vertex Number | Degree | Hub Score |
|---|---|---|---|---|---|
| Producer | Pinnularia | 1 | 35 | 6 | 0.5 |
| Producer | Achnanthidium | 1 | 36 | 2 | 0.24 |
| Producer | Brachysira | 1 | 38 | 2 | 0.24 |
| Producer | Surirella | 1 | 39 | 11 | 0.76 |
| Producer | Cyclotella | 1 | 40 | 2 | 0.19 |
| Producer | Didymosphenia | 1 | 41 | 3 | 0.16 |
| Producer | Melosira | 1 | 42 | 3 | 0.24 |
| Producer | Meridion | 1 | 43 | 4 | 0.33 |
| Producer | Eunotia | 1 | 44 | 6 | 0.53 |
| Producer | Tabularia | 1 | 45 | 2 | 0.24 |
| Invertebrate | Polypedilum | 1 | 46 | 31 | 0.19 |
| Invertebrate | Tanytarsus | 1 | 48 | 19 | 0.52 |
| Invertebrate | Procladius | 1 | 51 | 28 | 0.18 |
| Invertebrate | Micropsectra | 1 | 52 | 10 | 0.13 |
| Invertebrate | Chaetocladius | 1 | 53 | 3 | 0.13 |
| Invertebrate | Simulium | 1 | 55 | 21 | 0.37 |
| **Invertebrate** | **Cricotopus*** | 1 | 56 | **39** | 0.44 |
| Invertebrate | Conchapelopia | 1 | 58 | 5 | 0.13 |
| Invertebrate | Cryptochironomus | 1 | 59 | 15 | 0.13 |
| Producer | Planothidium | 1 | 60 | 6 | 0.51 |
| Producer | Sellaphora | 1 | 61 | 2 | 0.24 |
| Producer | Amphora | 1 | 62 | 14 | 0.88 |
| Invertebrate | Tipula | 1 | 63 | 12 | 0.02 |
| Invertebrate | Orthocladius | 1 | 64 | 10 | 0.27 |
| Invertebrate | Rheotanytarsus | 1 | 65 | 7 | 0.4 |
| **Invertebrate** | **Hydroptila*** | 1 | 66 | **39** | 0.13 |
| Invertebrate | Paratanytarsus | 1 | 67 | 6 | 0.13 |
| Invertebrate | Glyptotendipes | 1 | 68 | 5 | 0.24 |
| **Invertebrate** | **Enallagma*** | 1 | 69 | **33** | 0.18 |
| Invertebrate | Thienemanniella | 1 | 70 | 5 | 0.14 |
| Invertebrate | Parachironomus | 1 | 71 | 7 | 0.13 |
| Invertebrate | Baetis | 1 | 72 | 28 | 0.32 |
| Invertebrate | Bezzia | 1 | 73 | 8 | 0.21 |
| Invertebrate | Nais | 1 | 77 | 2 | 0.26 |
| Invertebrate | Stenelmis | 1 | 78 | 6 | 0.13 |
| Invertebrate | Lymnaea | 1 | 79 | 11 | 0.03 |
| Invertebrate | Ilyocryptus | 1 | 80 | 1 | 0.06 |
| Invertebrate | Eiseniella tetraedra | 1 | 81 | 10 | 0 |
| **Invertebrate** | **Hydropsyche*** | 1 | 82 | **48** | 0 |
| Invertebrate | Pisidium | 1 | 83 | 14 | 0 |

| Category | Taxon | Group | Vertex Number | Degree | Hub Score |
|---|---|---|---|---|---|
| Invertebrate | Eukiefferiella claripennis | 1 | 84 | 7 | 0 |
| Invertebrate | Simulium vittatum | 1 | 85 | 4 | 0 |
| Invertebrate | Diamesa | 1 | 86 | 13 | 0 |
| Invertebrate | Antocha saxicola | 1 | 87 | 1 | 0 |
| Invertebrate | Stenelmis crenata | 1 | 88 | 1 | 0 |
| Invertebrate | Sphaerium | 1 | 89 | 4 | 0 |
| Invertebrate | Libellula | 1 | 90 | 16 | 0 |
| Invertebrate | Zapada | 1 | 91 | 5 | 0 |
| Invertebrate | Rheocricotopus | 1 | 92 | 5 | 0 |
| Invertebrate | Gyraulus | 1 | 93 | 3 | 0 |
| Invertebrate | Prodiamesa | 1 | 94 | 4 | 0 |
| Invertebrate | Demicryptochironomus | 1 | 97 | 1 | 0 |
| Invertebrate | Admontia | 1 | 102 | 1 | 0 |
| Invertebrate | Chimarra | 1 | 103 | 1 | 0 |
| Invertebrate | Armadillidium | 2 | 49 | 3 | 0 |
| Invertebrate | Anelosimus | 2 | 100 | 1 | 0 |
| Invertebrate | Philoscia | 3 | 74 | 2 | 0 |
| Invertebrate | Dyschirius | 3 | 98 | 2 | 0 |
| Invertebrate | Lumbricus | 4 | 76 | 2 | 0 |
| Invertebrate | Carabus | 4 | 99 | 4 | 0 |
| Invertebrate | Chironomus | 5 | 47 | 3 | 0.08 |
| Invertebrate | Paramerina | 5 | 96 | 6 | 0 |
| Invertebrate | Fannia | 6 | 57 | 1 | 0 |
| Invertebrate | Hydrotaea | 6 | 101 | 1 | 0 |
| Producer | Ctenophora | 7 | 37 | 1 | 0 |
| Invertebrate | Colossendeis | 7 | 95 | 1 | 0 |
| Invertebrate | Drosophila | 8 | 50 | 2 | 0 |
| Invertebrate | Mesocyclops | 9 | 54 | 4 | 0 |
| Invertebrate | Microplitis | 10 | 75 | 2 | 0 |

* in the top 3 based on degree

** in the top 3 based on hub score

**Supplementary Table 7. Clair15 (good) clusters and keystone taxa (in bold).**

Members of the same group were given the same numeric designation.

| Category | Taxon | Group | Vertex Number | Degree | Hub Score |
|---|---|---|---|---|---|
| Producer | Cocconeis placentula | 1 | 8 | 8 | 0.92 |
| Producer | Melosira varians | 1 | 9 | 4 | 0.57 |
| Producer | Navicula cryptocephala | 1 | 11 | 2 | 0.37 |
| Producer | Navicula | 1 | 12 | 5 | 0.77 |
| Producer | Caloneis | 1 | 15 | 2 | 0.4 |
| Producer | Gomphonema | 1 | 16 | 5 | 0.77 |
| Producer | Encyonema | 1 | 17 | 3 | 0.41 |
| Producer | Cymbella | 1 | 18 | 6 | 0.84 |
| Producer | Nitzschia | 1 | 19 | 3 | 0.54 |
| **Producer** | **Rhoicosphenia**** | 1 | 20 | 8 | **0.99** |
| Producer | Pinnularia | 1 | 21 | 2 | 0.37 |
| Producer | Surirella | 1 | 24 | 5 | 0.65 |
| **Invertebrate** | **Polypedilum*** | 1 | 28 | **17** | 0.05 |
| Producer | Planothidium | 1 | 31 | 3 | 0.5 |
| **Producer** | **Amphora**** | 1 | 33 | 7 | **0.93** |
| **Invertebrate** | **Dugesia*** | 1 | 36 | **28** | 0 |
| **Invertebrate** | **Baetis*** | 1 | 37 | **18** | 0 |
| Invertebrate | Diamesa | 1 | 41 | 8 | 0 |
| Invertebrate | Tipula | 1 | 44 | 9 | 0 |
| Producer | Navicula gregaria | 2 | 1 | 4 | 0.61 |
| Producer | Navicula lanceolata | 2 | 2 | 4 | 0.61 |
| Producer | Amphora pediculus | 2 | 4 | 5 | 0.68 |
| Producer | Rhoicosphenia abbreviata | 2 | 6 | 6 | 0.78 |
| Producer | Navicula tripunctata | 2 | 10 | 4 | 0.61 |
| Invertebrate | Eiseniella tetraedra | 2 | 38 | 8 | 0 |
| Invertebrate | Pisidium | 2 | 39 | 12 | 0 |
| Invertebrate | Orthocladius | 2 | 40 | 6 | 0 |
| **Producer** | **Fragilaria**** | 3 | 5 | 10 | **1** |
| Producer | Frustulia | 3 | 13 | 3 | 0.42 |
| Producer | Achnanthes | 3 | 27 | 8 | 0.85 |
| Invertebrate | Stenelmis | 3 | 43 | 5 | 0 |
| Invertebrate | Rheocricotopus | 3 | 45 | 2 | 0 |
| Invertebrate | Dicranota | 3 | 46 | 4 | 0 |
| Invertebrate | Armadillidium | 4 | 29 | 1 | 0 |

| Category | Taxon | Group | Vertex Number | Degree | Hub Score |
|---|---|---|---|---|---|
| Invertebrate | Lumbricus | 4 | 34 | 2 | 0 |
| Invertebrate | Carabus | 4 | 48 | 3 | 0 |
| Producer | Diatoma vulgaris | 5 | 3 | 1 | 0.22 |
| Producer | Surirella brebissonii | 6 | 7 | 1 | 0.22 |
| Producer | Neidium | 7 | 14 | 1 | 0.22 |
| Producer | Achnanthidium | 8 | 22 | 1 | 0.22 |
| Producer | Brachysira | 9 | 23 | 1 | 0.22 |
| Producer | Gomphoneis | 10 | 25 | 1 | 0.14 |
| Producer | Meridion | 11 | 26 | 1 | 0.18 |
| Invertebrate | Drosophila | 12 | 30 | 2 | 0 |
| Producer | Sellaphora | 13 | 32 | 1 | 0.22 |
| Invertebrate | Deroceras | 14 | 35 | 2 | 0 |
| Invertebrate | Stenelmis crenata | 15 | 42 | 1 | 0 |
| Invertebrate | Paraleptophlebia | 16 | 47 | 1 | 0 |

* in the top 3 based on degree

** in the top 3 based on hub score

**Supplementary Table 8. Laurel7 (fair) clusters and keystone taxa (in bold).**

Members of the same group were given the same numeric designation.

| Category | Taxon | Group | Vertex Number | Degree | Hub Score |
|---|---|---|---|---|---|
| Invertebrate | Philoscia | 1 | 72 | 3 | 0 |
| Invertebrate | Bathyphantes | 1 | 95 | 1 | 0 |
| Invertebrate | Amara | 1 | 96 | 2 | 0 |
| Invertebrate | Carabus | 1 | 97 | 2 | 0 |
| Producer | Staurosira construens | 2 | 1 | 2 | 0.25 |
| Producer | Navicula gregaria | 2 | 2 | 8 | 0.68 |
| Producer | Navicula lanceolata | 2 | 3 | 7 | 0.64 |
| Producer | Diatoma | 2 | 4 | 13 | 0.89 |
| Producer | Diatoma vulgaris | 2 | 5 | 2 | 0.25 |
| Producer | Meridion circulare | 2 | 6 | 8 | 0.53 |
| Producer | Amphora pediculus | 2 | 7 | 9 | 0.71 |
| **Producer** | **Fragilaria**** | 2 | 8 | 17 | **1** |
| Producer | Gomphonema truncatum | 2 | 9 | 5 | 0.49 |
| Producer | Gomphonema acuminatum | 2 | 10 | 4 | 0.33 |
| Producer | Epithemia | 2 | 11 | 7 | 0.42 |
| Producer | Sellaphora seminulum | 2 | 12 | 2 | 0.25 |
| Producer | Amphora ovalis | 2 | 13 | 8 | 0.61 |
| Producer | Nitzschia linearis | 2 | 14 | 1 | 0.06 |
| Producer | Navicula cryptotenella | 2 | 15 | 2 | 0.25 |
| Producer | Cymbella | 2 | 16 | 11 | 0.87 |
| Producer | Nitzschia amphibia | 2 | 17 | 2 | 0.25 |
| Producer | Surirella brebissonii | 2 | 18 | 3 | 0.35 |
| Producer | Nitzschia dissipata | 2 | 19 | 9 | 0.76 |
| Producer | Cocconeis placentula | 2 | 20 | 14 | 0.84 |
| Producer | Melosira varians | 2 | 21 | 9 | 0.76 |
| Producer | Navicula tripunctata | 2 | 22 | 8 | 0.68 |
| Producer | Cymbella tumida | 2 | 23 | 1 | 0.06 |
| Producer | Navicula capitatoradiata | 2 | 24 | 2 | 0.25 |
| Producer | Eunotia bilunaris | 2 | 25 | 2 | 0.25 |
| Producer | Navicula cryptocephala | 2 | 27 | 4 | 0.43 |
| Producer | Staurosira | 2 | 28 | 9 | 0.77 |
| Producer | Navicula | 2 | 29 | 11 | 0.91 |
| Producer | Frustulia | 2 | 30 | 8 | 0.72 |
| Producer | Neidium | 2 | 31 | 4 | 0.45 |

| Category | Taxon | Group | Vertex Number | Degree | Hub Score |
|---|---|---|---|---|---|
| Producer | Caloneis | 2 | 32 | 5 | 0.5 |
| **Producer** | **Gomphonema**** | 2 | 33 | 12 | **0.94** |
| Producer | Encyonema | 2 | 34 | 7 | 0.68 |
| Producer | Nitzschia | 2 | 35 | 6 | 0.56 |
| **Producer** | **Rhoicosphenia**** | 2 | 36 | 15 | **1** |
| Producer | Pinnularia | 2 | 37 | 8 | 0.7 |
| Producer | Achnanthidium | 2 | 38 | 4 | 0.45 |
| Producer | Brachysira | 2 | 39 | 4 | 0.45 |
| Producer | Surirella | 2 | 40 | 13 | 0.87 |
| Producer | Psammothidium | 2 | 41 | 7 | 0.68 |
| Producer | Synedra | 2 | 43 | 13 | 0.78 |
| Producer | Meridion | 2 | 44 | 2 | 0.18 |
| Producer | Gyrosigma | 2 | 45 | 2 | 0.1 |
| Producer | Cocconeis | 2 | 46 | 4 | 0.34 |
| Producer | Tabularia | 2 | 47 | 4 | 0.45 |
| Invertebrate | Polypedilum | 2 | 48 | 27 | 0.01 |
| Invertebrate | Tanytarsus | 2 | 50 | 16 | 0.26 |
| Invertebrate | Micropsectra | 2 | 52 | 10 | 0 |
| Invertebrate | Microtendipes | 2 | 53 | 21 | 0 |
| Invertebrate | Simulium | 2 | 55 | 24 | 0.31 |
| Invertebrate | Cricotopus | 2 | 57 | 39 | 0.32 |
| Invertebrate | Conchapelopia | 2 | 59 | 4 | 0.12 |
| Invertebrate | Cryptochironomus | 2 | 60 | 12 | 0 |
| Producer | Planothidium | 2 | 61 | 7 | 0.68 |
| Producer | Sellaphora | 2 | 62 | 4 | 0.45 |
| **Producer** | **Amphora**** | 2 | 63 | 13 | **0.93** |
| Invertebrate | Tipula | 2 | 64 | 14 | 0 |
| Invertebrate | Orthocladius | 2 | 65 | 13 | 0.27 |
| **Invertebrate** | **Hydroptila*** | 2 | 66 | **44** | 0.15 |
| Invertebrate | Paratanytarsus | 2 | 67 | 4 | 0 |
| Invertebrate | Caenis | 2 | 69 | 24 | 0.01 |
| Invertebrate | Thienemanniella | 2 | 70 | 7 | 0.14 |
| Invertebrate | Parachironomus | 2 | 71 | 3 | 0 |
| Invertebrate | Pisidium | 2 | 75 | 15 | 0.01 |
| **Invertebrate** | **Dugesia*** | 2 | 77 | **42** | 0 |
| Invertebrate | Eiseniella tetraedra | 2 | 78 | 9 | 0 |
| **Invertebrate** | **Hydropsyche*** | 2 | 79 | **43** | 0 |
| Invertebrate | Simulium vittatum | 2 | 80 | 5 | 0 |
| Invertebrate | Dicrotendipes | 2 | 81 | 3 | 0 |
| Invertebrate | Antocha saxicola | 2 | 82 | 1 | 0 |

| Category | Taxon | Group | Vertex Number | Degree | Hub Score |
| --- | --- | --- | --- | --- | --- |
| Invertebrate | Stenelmis crenata | 2 | 83 | 1 | 0 |
| Invertebrate | Sphaerium | 2 | 84 | 4 | 0 |
| Invertebrate | Paraleptophlebia | 2 | 85 | 1 | 0 |
| Invertebrate | Libellula | 2 | 89 | 12 | 0 |
| Invertebrate | Oecetis | 2 | 91 | 3 | 0 |
| Invertebrate | Camponotus | 2 | 92 | 1 | 0 |
| Invertebrate | Orconectes | 2 | 93 | 1 | 0 |
| Invertebrate | Rhyacophila | 2 | 94 | 10 | 0 |
| Invertebrate | Nais | 3 | 76 | 3 | 0.24 |
| Invertebrate | Stenophylax | 3 | 100 | 1 | 0 |
| Invertebrate | Conocephalus | 4 | 73 | 2 | 0 |
| Invertebrate | Argiope | 4 | 98 | 2 | 0 |
| Invertebrate | Sitona | 5 | 74 | 4 | 0 |
| Invertebrate | Tachinus | 5 | 99 | 1 | 0 |
| Invertebrate | Chironomus | 6 | 49 | 2 | 0.01 |
| Invertebrate | Paramerina | 6 | 90 | 4 | 0 |
| Producer | Cyclotella | 7 | 42 | 3 | 0.19 |
| Invertebrate | Daphnia | 7 | 86 | 2 | 0 |
| Producer | Navicula radiosa | 8 | 26 | 1 | 0.03 |
| Invertebrate | Drosophila | 9 | 51 | 2 | 0 |
| Invertebrate | Mesocyclops | 10 | 54 | 4 | 0 |
| Invertebrate | Stagmomantis | 11 | 56 | 4 | 0 |
| Invertebrate | Nanocladius | 12 | 58 | 1 | 0 |
| Invertebrate | Glyptotendipes | 13 | 68 | 1 | 0 |
| Invertebrate | Stempellinella | 14 | 87 | 1 | 0 |
| Invertebrate | Metriocnemus | 15 | 88 | 1 | 0 |

* in the top 3 based on degree

** in the top 3 based on hub score

**Supplementary Figure 1. Sampling sites and condition.** Terrain map tiles by Stamen Design, under CC BY 3.0. Data by OpenStreetMap, under ODbL.

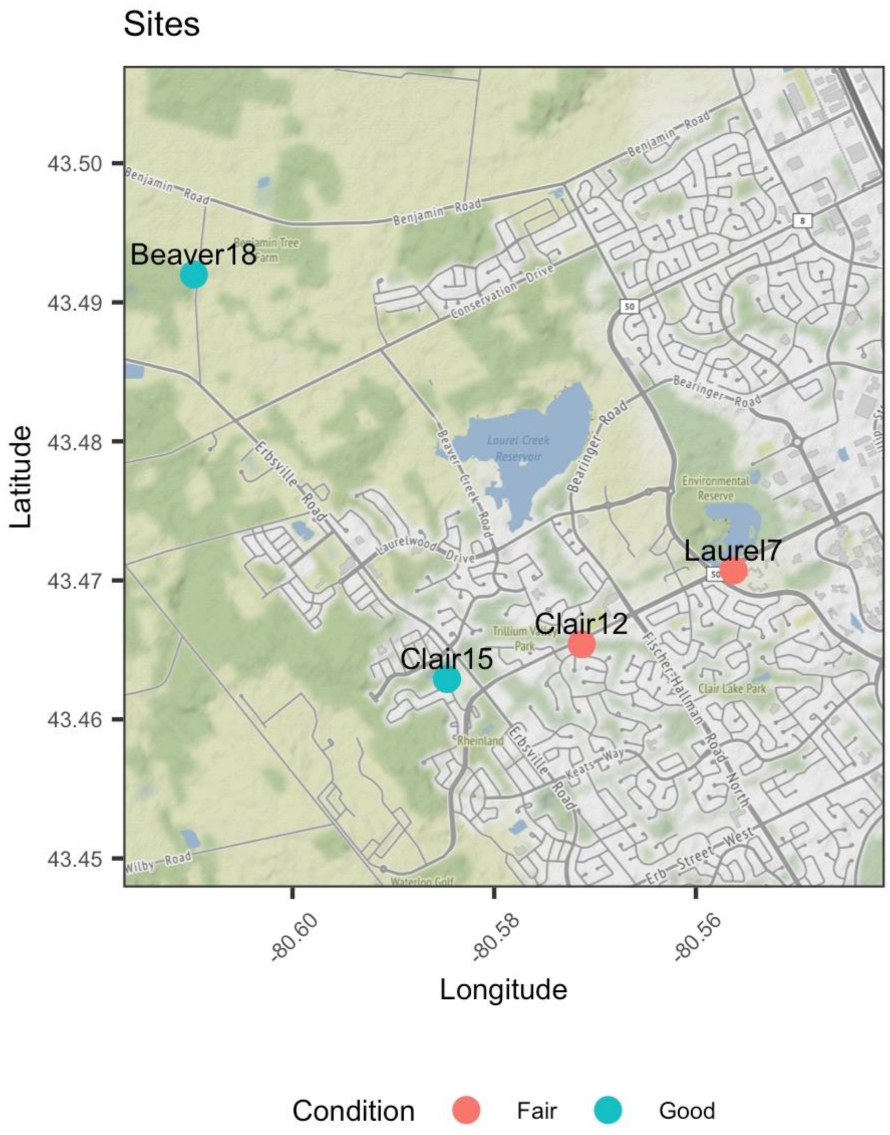

**Supplementary Figure 2. All samples show that ESV sampling reached saturation.** Each line represents reads from a sample plotted against the number of detected ESVs.  Samples were color-coded by site (Beaver 18, Clair 12, Clair 15 and Laurel7), marker (rbcL and COI), or site status as shown in the legends.  The vertical dashed line indicates the 15th percentile of sampling read depth (45,937), which is the number of macroinvertebrates and diatoms reads that would be accounted for in any ESV level analysis based on normalized data.

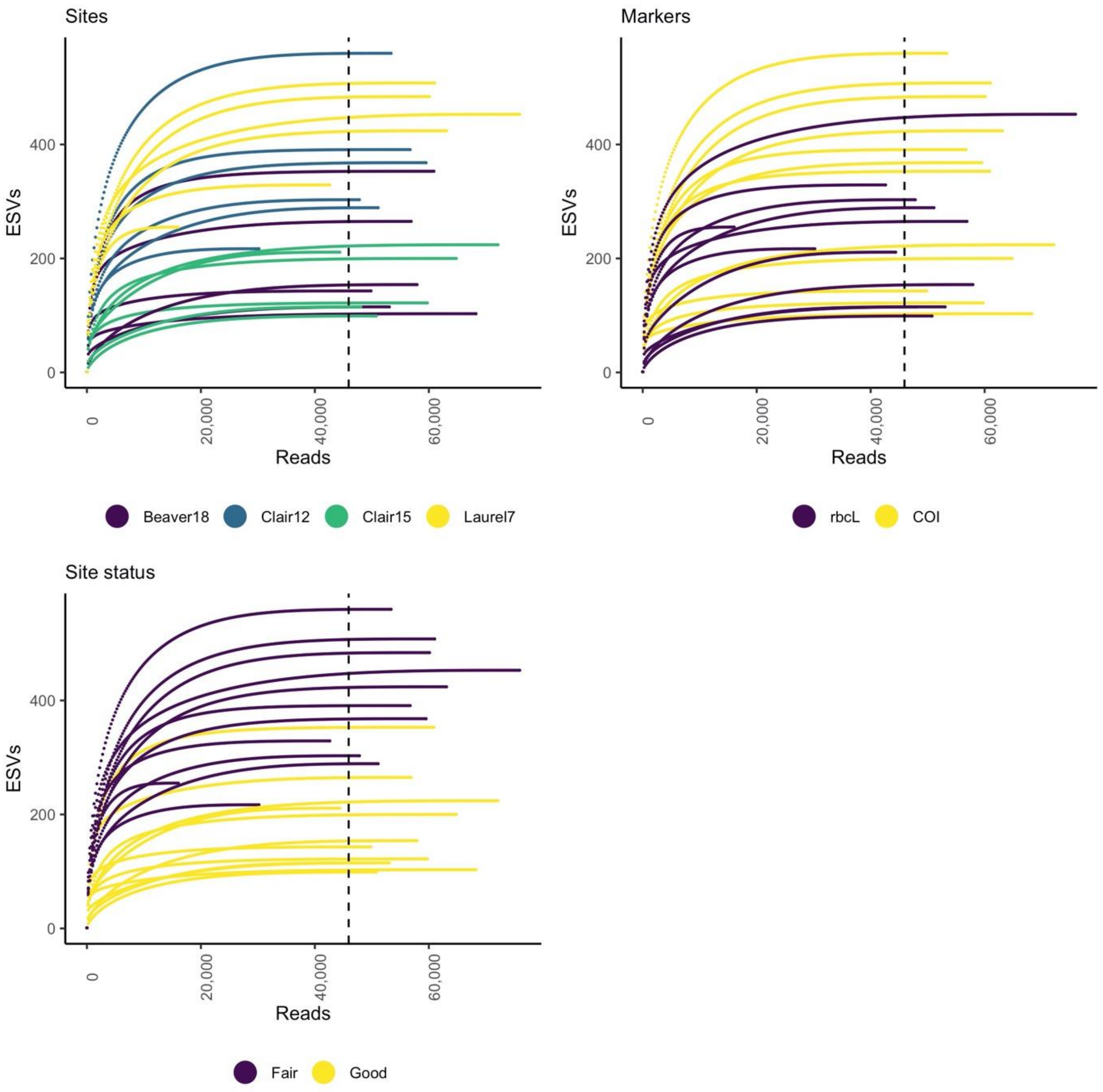

**Supplementary Figure 3. Diatom community composition differs between fair and good sites.** Results are summarized to the order rank to facilitate readability. Reads are shown on a log10 scale. Labels on the x axis represent field replicates, sites, site condition, and number of orders detected. Based on normalized data.

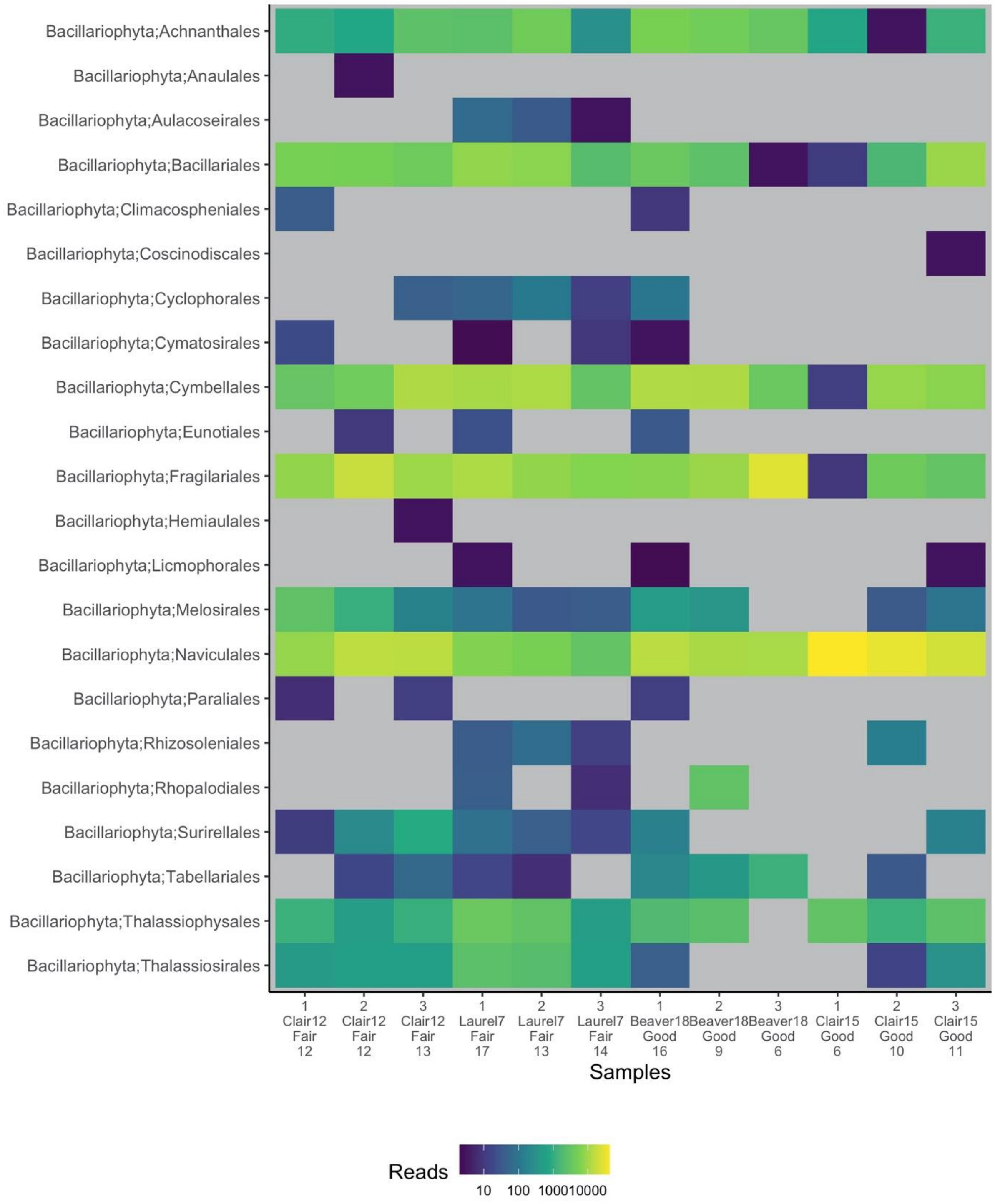

**Supplementary Figure 4. Macroinvertebrate community composition differs between fair and good sites.** Results are summarized to the order rank to facilitate readability. Reads are shown on a log10 scale. Labels on the x axis represent field replicates, sites, site condition, and number of orders detected. Based on normalized data.

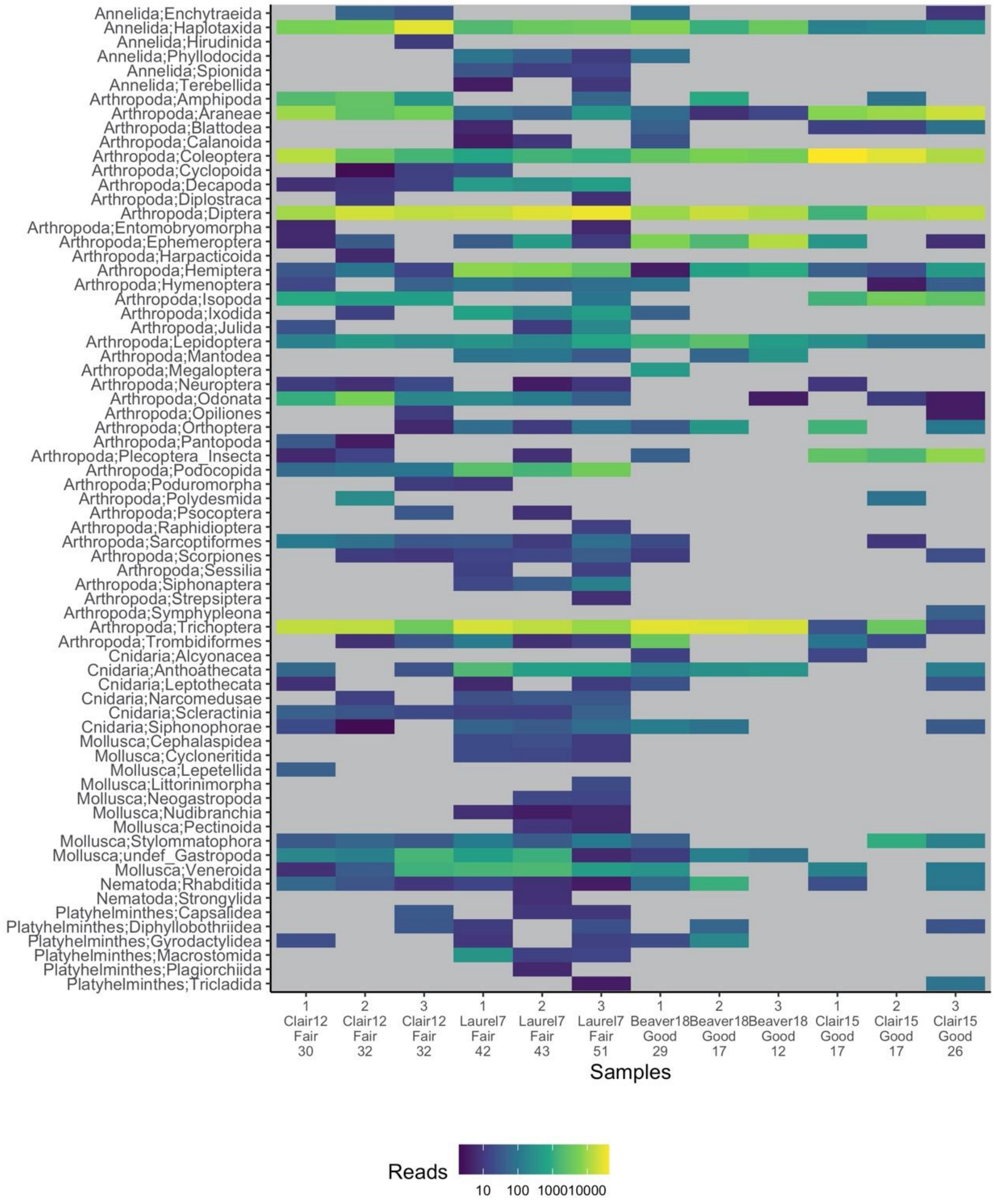

**Supplementary Figure 5. Pairwise community dissimilarities within site condition groupings tended to be larger among 'good' sites for diatoms.** The distribution of within-group pairwise community dissimilarities for macroinvertebrates was largely overlapping, skewing slightly larger for 'good' sites. Based on binary Bray Curtis dissimilarities where read counts were initially normalized using rarefaction to the 15th percentile.

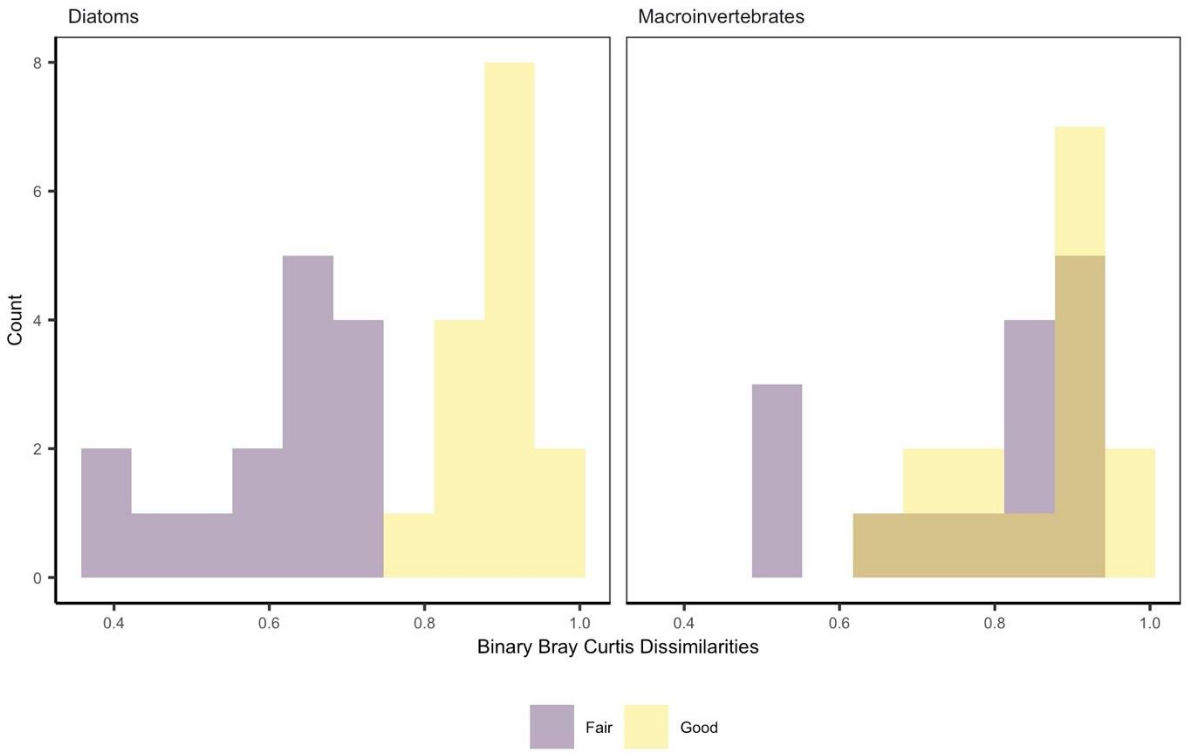

**Supplementary Figure 6. A suit of site status indicators were detected using two methods.** Indicator species were detected using the indicator value method (IndVal.g) and plotted along with the A (specificity or positive predictive value) and B (fidelity or sensitivity) components; as well as using the point biserial correlation coefficient (r.g). Only indicators with a p-value < 0.05 are shown. For each statistic, the value ranges from 0 to 1, where the higher the value, the better the indicator. We show A) macroinvertebrate and B) diatom site status indicators. Results based on a species x sample matrix with rarefied read counts. The cnidarian taxon below is likely misidentified due to ‘over classification’, i.e., the taxonomic assignment had low bootstrap support values and likely represents a taxon not present in the underlying reference database and was removed from the analyses presented in the main text.

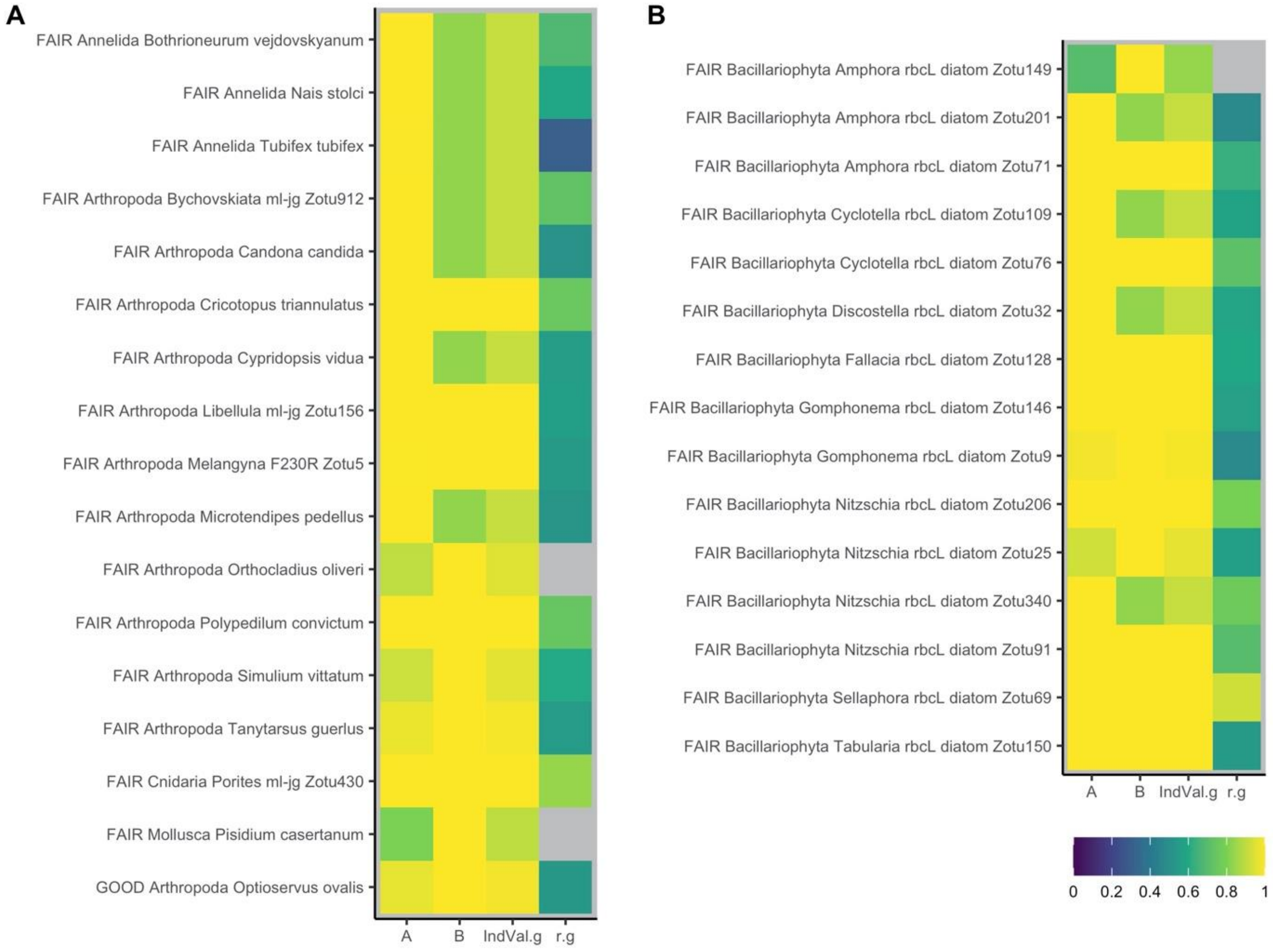
